# Next-generation all-in-one CRISPR/Cas9 multiply-edited CD30CAR-T cells: Potency despite risk of translocations

**DOI:** 10.64898/2026.02.27.708432

**Authors:** Jan-Malte Kleid, Michael Damrat, Milita Dargužytė, Manuel Rhiel, Natascha E. Stumpf, Tabea Kleitke, Sandra Ammann, Tatjana I. Cornu, Fawad Khan, Tobias Wollmann, Sven Borchmann, Christof Scheid, Christianne Moraes, Tobias Riet, Sabine Awerkiew, Leon Ullrich, Birgit Gathof, Frank Klawonn, Britta Eiz-Vesper, Dimitrios L. Wagner, Kai Hübel, Roland Ullrich, Martin Bornhäuser, Toni Cathomen, Renata Stripecke

## Abstract

**Background:** Chimeric antigen receptor (CAR)-T cells are therapeutic breakthroughs against advanced non-Hodgkin lymphomas and myelomas. On the other hand, no CAR-T cell product has been so far clinically approved for therapy of Hodgkin Lymphoma (HL), T cell lymphoma (TCL), or Epstein-Barr-Virus (EBV)-associated lymphoproliferative diseases (EBV-LPDs). CD30 (TNFRSF8) is commonly expressed on HL and on subsets of TCL and EBV-LPDs. CD30CAR-T cells generated via transduction with viral vectors have been tested in clinical trials, showing overall good responses against HL. CAR-T cells produced entirely with locus-specific gene editing methods are emerging as attractive next-generation engineered cell products for ease of multiple seamless cell modifications.

**Methods:** Using CRISPR/Cas9-mediated techniques, we optimized homology-directed repair templates (HDRTs) and performed all-in-one multiplex editing to knock-in (KI) CD30CAR within the TCRα constant (*TRAC*) locus and to simultaneously knock-out (KO) PD-1 or/and β2M. CD30CAR-T cells were tested in CD30^+^ cell models of HL, TCL, and EBV-LPDs.

**Results:** We compared mouse versus human anti-CD30 scFv designs in HDRTs incorporating *TRAC* homology arms, FcIg spacer/detection domain, and CD28 / CD3ζ signaling domains. We obtained an average of 30% TRAC^KI^CD30CAR-T cells and efficient *in vitro* cytotoxicity with CD30^+^ cell targets. CARs incorporating the high-affinity humanized 5F11 scFv showed the highest CAR expression, and the editing templates were further modified to incorporate a truncated CD34 (tCD34) spacer/detection domain. 5F11-CD30CAR-tCD34-T cells showed high CAR-KI rates (approx. 50-80% 12-14 days after editing) and potency *in vitro* and *in vivo*. Subsequently, we tested all-in-one CAR KI with additional KOs by co-electroporation of guide RNAs (gRNAs) targeting the genes encoding PD-1 or /and β2M to improve function and allow for improved cell persistence in allogeneic recipients, respectively. Compared with CD30CAR-T cells, CD30CAR-β2M^KO^-T cells were similarly viable and functional and showed low risk of translocations. PD1^KO^ enabled CD30CAR-T cells to produce higher levels of cytotoxic features upon exposure to targets. However, simultaneous β2M^KO^ and PD-1^KO^ compromised the expansion capacity of CD30CAR-T cells and resulted in detectable translocations.

**Conclusions:** Non-virally engineered 5F11-CD30CAR-T cells represent a novel cell therapy modality against CD30^+^ lymphomas. Multiplex editing remains to be optimized to avoid unwanted genomic alterations and chromosomal translocations.

## Background

CD30 (TNFRSF8) is a member of the tumor necrosis factor receptor superfamily. CD30 limits the proliferation of autoreactive CD8^+^ T cells and is expressed in a subset of memory T and B cells [1]. Previous studies have identified the pro-oncogenic roles of CD30 in classical Hodgkin lymphoma (cHL), anaplastic large cell lymphoma (ALCL), and a portion of diffuse large B-cell lymphoma (DLBCL) cases [2]. CD30 upregulation in lymphomagenesis was associated with anti-apoptotic mechanisms and cell survival [1]. Recent studies have found correlations between CD30 upregulation and viral infection and reactivations. For example, there is a direct cause-effect relationship between EBV infection, expression of the viral oncoprotein LMP1 with oncogenic effects, and upregulation of CD30 expression [3]. Moreover, CD30 is upregulated in EBV-positive lymphomas, as well as on polyclonal B cells infected with EBV [1]. EBV^pos^ DLBCL has a higher percentage of cells expressing CD30 and is associated with lower survival than EBV^neg^ DLBCL [4].

Chimeric antigen receptor (CAR)-reprogrammed immune cells have emerged as interesting novel therapeutic tools against cancer, autoimmune diseases, and infections. Abken and colleagues were pioneers in the development of retrovirally engineered CD30CAR-T cells incorporating the HRS-3 mouse scFv monoclonal antibody sequence targeting CD30 and demonstrated that HRS-3-CD30CAR-T cells showed high and specific cytotoxicity against cHL [5,6]. Led by Heslop, Savoldo and colleagues, HRS-3/CD28ζ-CD30CAR-T cells generated with retroviral vectors and administered after fludarabine-based lymphodepletion were extensively tested clinically against cHL [7–9]. These HRS-3/CD28ζ-CD30CAR-T cells showed a favorable safety profile and excellent anti-tumor responses in heavily pretreated patients with relapsed/refractory (r/r) cHL (NCT#04268706, phase 1b/2 trial). Doses of HRS-3/CD28ζ-CD30CAR-T cells were also tested as consolidation therapy after hematopoietic stem cell transplantation of high-risk CD30^pos^ cHL and Non-Hodgkin lymphoma (NHL) patients, showing feasibility and low occurrences of cytokine release syndrome (NCT#02663297 phase 1 trial) [10]. Briones and colleagues developed CAR-T cells targeting a proximal epitope within the CD30 molecule and used enriched memory T cells for lentiviral transductions to ensure persistence [11]. CD30CAR-T treatment of r/r cHL and CD30^pos^ T-cell non-Hodgkin lymphoma (T-NHL) was well tolerated, and there were no dose-limiting toxicities (DLTs) (NCT#04653649). Kochenderfer and colleagues reported a phase 1 dose escalation trial of CD30CAR-T cells manufactured with lentiviral vectors and incorporating CD28ζ−signaling domains and scFvs derived from the high-affinity humanized 5F11 monoclonal antibody [12]. The patients enrolled had a high tumor burden and were heavily pretreated (average of 7 prior lines of therapy). The trial showed an overall response rate of 43% but was halted due to substantial toxicities (NCT#03049449) [12]. Overall, the few currently available long-term CD30CAR-T clinical data suggest a shorter duration of responses compared with the vast experience with CD19CAR-T cells. Nevertheless, given the urgent need for high-risk r/r CD30^+^ lymphoma cases, new CD30CAR-T production modalities are needed to improve their potency, persistence, safety, and availability.

CAR-T cells have been mostly engineered with γ-retroviral vectors and lentiviral vectors, which are classified at biosafety level 2 and randomly integrate in the genome. Although the clinical safety profile has been shown to be favorable, these non-specific gene insertions may cause clonal expansion, insertional mutagenesis, and variable transgene expression. Non-viral clustered regularly interspaced short palindromic repeats (CRISPR)/Cas technology combined with homology-directed repair templates (HDRTs) are based on DNA or RNA, hence with lower biosafety concerns, and can precisely integrate within genomic loci in T cells [13]. Sadelain, Eyquem, and colleagues were the first to show that introducing CD19CAR into the T-cell receptor (TCR) α constant (*TRAC*) locus via gene editing resulted in uniform CAR expression in human peripheral blood T cells and notably enhanced *in vivo* T-cell potency compared with CD19CAR-T cells generated with viral vectors causing random integrations [14]. The interpretation of these superior results inferred that targeting the CAR to the *TRAC* locus provided a physiological modulation of the CAR signaling and expression of the CAR following exposures to antigen for better control of effector T-cell differentiation and reduction of T-cell exhaustion [14]. Recent advances in CRISPR/Cas technology with efficient HDR, promoted site-directed non-viral CAR-target gene knock-in (KI) into the *TRAC* locus, resulting in up to 50% TRAC^KI^CAR-T cells [13,15,16]. CD19CAR inserted into the *TRAC* locus via CRISPR/Cas-mediated homology-directed repair simultaneously prevents TCR expression, reducing interferences of TCR and CAR signaling. We also observed that compared with LV-CD19CAR-T cells, gene-edited TRAC^KI^CD19CAR-T cells showed improved *in vivo* performance and persistence against human Nalm-6 leukemia growth in xenografted mouse models [15].

Here, we sought to advance towards the generation of TRAC^KI^CD30CAR-T cells. We systematically tested new scFv binders and spacers/detection domains incorporated in the HDRT templates. Interestingly, expanded TRAC^KI^CD30CAR-T cells showed high frequencies of naïve and central memory T cell phenotypes. CD30CAR-T cells killed HL cells *in vitro* and *in vivo*, showing high persistency and potential for clinical development. Multiple all-in-one CRISPR-mediated knockouts to abolish expression of programmed cell death protein 1 (PD)-1 and β2-microglobulin (β2M) were feasible. CD30CAR-T cells with PD-1^KO^ killed different lymphoma cell types expressing PD-L1 and upregulated the expression of cytotoxic proteins but displayed genomic rearrangements in the form of translocations. Our studies advance towards new strategies to generate non-viral CD30CAR-T cells with new features, such as lacking expression of β2M/ HLA-I for the generation of partially matched allogeneic products.

## Methods

### Cell lines

L540 (ACC72) and L591 (ACC602) cell lines were obtained from the German Collection of Microorganisms and Cell Cultures (DSMZ, Brunswick, Germany). Myla cells were kindly provided by Dr. T. Braun (University Hospital Cologne, Cologne, Germany). Lymphoblastoid cell lines (LCLs) infected with EBV were produced in R. Stripecke’s previous laboratory at the Hannover Medical School (Hannover, Germany) [17]. Lymphoma cell lines were passaged in RPMI 1640 (Gibco^TM^ by Thermo Fisher Scientific, Waltham, MA) with 10 % FBS and 1x P/S. All cell cultures were maintained at 37°C and 5 % CO_2_. Cell lines expressing fluc-GFP were generated after transduction with the lentiviral vector pRRL.PPT.Cbx3-SFFV-fLuc-T2A-eGFP (the plasmid DNA containing the backbone vector was kindly provided by Prof. Axel Schambach, Institute of Experimental Hematology, Hannover Medical School, Hannover, Germany). The GFP^bright^ cells were sorted, pools and clonal lines were expanded, and stocks were subsequently cryopreserved at −150°C.

### Quantification of EBV copies

DNA isolation of mouse BM to determine the EBV load in target cell lines was performed with a QIAamp DNA Blood Mini Kit (Qiagen, Valencia, CA) according to the manufacturer’s protocol. DNA concentration and purity were measured using the Nanodrop instrument. EBV DNA detection by real-time PCR was performed with the EBV in vitro diagnostics PCR kit (GeneProof®, Dolni Herspice, Brno, Czechia) after amplification of a specific conserved DNA sequence of the single-copy gene encoding the nuclear antigen 1 (EBNA1) and measurement of fluorescence emission. An internal standard (IS) was included in the reaction mix as a control for the PCR reaction. The detection limit of the assay was approximately 1.9 × 10^5^ EBV units in the sample.

### Design, amplification and purification of HDRTs

The CD19CAR HDRT sequence with the *TRAC* homology arms, an optimized signal peptide, the FMC63 CD19 scFv, the IgG1-CH3 FC detection domain, and the CD28/CD3ζ signaling domain was previously described [16]. The CD30CAR HDRTs were designed to incorporate the HRS3 scFv sequence [5,6] or humanized sequences of monoclonal antibodies binding CD30 (5F11 or 2H9 described in the patent application (US7387776B2 - Human monoclonal antibodies against CD30) [18]. The sequence of the tCD34 detection domain was adapted from a previous publication [19]. The plasmids containing HDRT sequences were synthesized by Twist Bioscience (San Francisco, CA). HDRTs were amplified by PCR using Herculase II Fusion DNA Polymerase (Agilent, Santa Clara, CA) The *TRAC* HDR arms (forward 5′-ATAAAAGAATAAGCAGTATTATTAAGTAGCCCTGC-3’; reverse 5′- CATCTGCTTTTTTCC-CGTGTCATTCT −3’) were previously validated [15]. The PCR products were purified with DNA-binding magnetic beads (AMPure XP, Beckman Coulter, Brea, CA) and recovered with a DynaMag™-2 magnet (Thermo Fisher Scientific, Waltham, MA), washed with 70% ethanol, air dried, diluted in water and the concentrations of the HDRTs were measured with a Nanophotometer® (Implen, Munich, Germany). The HDRT DNAs were adjusted to a concentration of 1 µg/µl with nuclease-free water and cryopreserved at −20°C.

### Generation of gene-edited CAR-T cells

Peripheral blood mononuclear cells (PBMCs) were purified by density-gradient centrifugation with Pancoll (PAN Biotech, Aidenbach, Germany) and used immediately or cryopreserved. Prior to activation, PBMCs were pre-cultured for 2-3 days in RPMI with 10 % FCS and 1x P/S, supplemented with interleukin (IL)-7 (5 ng/ml) and IL-15 (5 ng/ml) (both from Miltenyi Biotec, Bergisch Gladbach, Germany) to reduce the presence of unwanted B cells or monocytes. PBMCs were activated with TransAct (Miltenyi Biotec) for 2 days using 10 µL TransAct per mL RPMI (10 µL/mL) for 2×10^6^ cells with 10% FBS and 1x P/S, supplemented with IL-7 and IL-15 (5 ng/ml each). Cells were washed, counted, and resuspended at 1×10^8^ cells/ml TexMACS (Miltenyi Biotec). 50 µl containing up to 5×10^6^ cells or 1 ml containing up to 1×10^8^ cells were used for small-scale (SS) or for medium-scale (MS) electroporation, respectively. To generate the RNP complex, crRNA and tracrRNA diluted in nuclease-free duplex buffer (IDT, Newark, NJ) were mixed and heated at 95 °C for 5 min to form the guide RNA. Dependent on the experiment setup, crRNAs targeting *TRAC* (5′-GAGAATCAAAATCGGTGAAT-3′), β2M (5′-CTTACCCCACTTAACTATCT-3′) or *PDCD1* (5′-CACCTACCTAAGAACCATCC-3′) were added. After cooling to RT, 15-50 kDa poly-L-glutamic acid (PGA) (Sigma Aldrich, St. Louis, MO) and 1 µM Cas9 (Alt-R S.p Cas9 Nuclease V3, 61 µM, IDT) were added, and the mixture was incubated for 30 min to allow RNP formation prior to adding the HDRT. R-50×8 cuvettes (ER050U8-03, MaxCyte, Gaithersburg, MD) and R-1000 cuvettes (ER001M1-10, MaxCyte) were used for SS and MS productions, respectively. An ExPERT ATx^TM^ Electroporation® device (MaxCyte) was used for electroporation with the “Resting T cell 14-3” program. After electroporation, the cells were cultured in TexMACS medium (Milteny Biotec) containing 2 µM HDR Enhancer V2 (IDT) for 24 h at 37°C. After 2 h, cytokines (IL-7 and IL-15, each 5 ng/ml) were added. After 24 h the cells were washed and cultured in RPMI + 10% FBS. Cytokines (IL-7 and IL-15, 5 ng/ml each) were added every 2-3 days. Cells were counted and analyzed by flow cytometry at several timepoints after electroporation and up to 9-14 days. For ease of conformity to test cells generated from different donors in potency assays, expanded CAR-T cells were cryopreserved in freezing medium (10% DMSO + 5% Glucose 20% human serum albumin in final volume diluted in PBS) and maintained at −150°C until use. Cryopreserved cells were thawed and cultured in R10 plus cytokines for 2 h prior to use for potency assays or *in vivo*. Lots of cryopreserved CD30CAR-T cells used for *in vivo* experiments were pre-tested for mycoplasma contamination.

### Flow cytometry analyses of gene-edited CAR-T cells and cell lines

As a first step, cells were blocked in 100 µl PBS containing 10 µg/ml mouse-IgG (Sigma-Aldrich, St. Louis, Missouri) and CAR-T cells were analyzed for CD3^KO^, CAR^KI^, viability, and expression of CD4 and CD8. For CAR-T cells with the FcIg-based spacer domain, CAR^KI^ was detected by staining with 1:100 Alexa Fluor® 647 “AffiniPure F(ab’)₂ Fragment Goat Anti-Human IgG” (109-606-170, Jackson Immuno Research Laboratories, West Grove, PA). The staining of the CAR with the anti-human IgG was done separately from the other subsequently used staining monoclonal antibodies (mAbs) to avoid complex formation. Staining for detection of CAR-T cells with tCD34 detection domain was performed with the antibody BD Pharmingen™ Alexa Fluor™ 647 Mouse Anti-Human CD34 (568771, BD Biosciences, Franklin Lakes, NJ). Staining for immunophenotype characterization of naïve, central memory, effector memory, and terminal effector cells was performed with antibodies detecting CD45RA (Brilliant Violet 605™ anti-human CD45RA, 304134, BioLegend, San Diego, CA) and CD62L (PE/Cyanine7 anti-human CD62L, 304822, BioLegend). Staining for detection of HLA-I knockout was performed with an antibody detecting residues in the N terminus of the human β2-microglobulin molecule (APC/Cyanine7 anti-human HLA-A,B,C, 311426, BioLegend). Detection of CD30 and PD-L1 on target cells was performed with commercially available antibodies (see supplement **Table S1** describing the antibodies and dilutions used for staining). The cells were stained for 30 min at 4°C in the dark, washed in PBS containing 1% FBS, resuspended in 100 µl PBS containing 1% FBS, and then analyzed using a Cytoflex S instrument (Beckman Coulter, Brea, CA).

### *In vitro* co-culture of CAR-T cells with target cells and bio-luminescence assays

For cytotoxic assays, target cells L540/fluc-GFP (clone E after sorting), L591/fluc-GFP (sorted pool), Myla/fluc-GFP (sorted pool), and LCLs/fluc-GFP (pool) were seeded in 96-well round-bottom plates (1×10^4^ cells per well) in 50 µL RPMI with 10 % FBS and 1x P/S supplemented with IL-7 and IL-15 (both 5 ng/ml). Total T cells recovered from the expanded cultures were added in the same medium at E:T ratios of 0.03: 1, 0.1: 1, 0.3:1, 1:1, 3:1 and 10:1 in 50 µL per well. All co-cultures were performed as technical triplicates and were incubated for 72 h (37°C and 5% CO_2_). After incubation, the cells were spun down, washed, and transferred into a white 96-well half-area plate. The cell pellets were resuspended in 100 µl D-luciferin potassium salt solution (2.5 mg in 1 ml PBS purchased from SYNCHEM, Felsberg, Germany). Luminescence emitted by living cells was measured for each triplicate with a TECAN Spark^®^ with injector module (TECAN, Männedorf, Switzerland).

### Testing CD30CAR-T cells in an *in vivo* xenograft model

All animal experiments have been approved by LAVE (Landesamt für Verbraucherschutz und Ernährung) approval number Az 81-02.04.2023.A155. 8-week-old female NSG-MHC I/II DKO mice (*NOD.Cg-Prkdc^scid^ H2-K1^b-tm1Bpe^ H2-Ab1^g7-em1Mvw^ H2-D1^b-tm1Bpe^ IL-2rg^tm1Wjl^/SzJ*) were obtained from the Jackson Laboratory (JAX; Bar Harbor, ME) and maintained under pathogen-free conditions. 1×10^6^ L540/fLuc-GFP cells diluted in 100 µl PBS were injected intravenously (i.v.) into the tail vein, and control mice were injected with 100 µl PBS. The first baseline BLI analyses to confirm tumor engraftment were performed on day 4 after lymphoma injection with the IVIS Spectrum apparatus (PerkinElmer, Waltham, MA) and the data were analyzed using LivingImage software (PerkinElmer) and Aura (Spectral Instruments) as previously described [15]. After anesthesia with isoflurane, mice were injected intraperitoneally (i.p.) with 2.5 mg D-Luciferin potassium salt dissolved in 100 µl PBS, and frontal and lateral images were obtained with an exposure time adjusted based on the intensity of the luminescence signal. On day 5 after tumor challenge, 1×10^6^ T cells diluted in 100 µl PBS were injected i.v. in CAR-T cell-treated mice or 100 µl PBS in tumor-only mice. During the experimental period, mice were monitored every second day regarding weight and signs of graft-versus-host disease (GvHD) or other distress, and mice showing signs of distress were monitored daily. BLI analyses after T cell infusion were performed biweekly up to 14 weeks after lymphoma challenge. After euthanasia, tissues were collected for FACS analyses. Macroscopic examinations of spleen (SPL), liver, and kidneys were performed. SPL were smashed through a 100 µm cell strainer to obtain single-cell suspensions. Bone marrow (BM) was flushed from the femora and tibia. Peripheral blood (PBL) and SPL samples were resuspended in RBC lysis buffer (Invitrogen, Carlsbad, USA) and incubated for 5 min at RT and then washed with PBS. The cells were used directly for FACS analysis, and leftover cells were cryopreserved in freezing medium (10 % DMSO + 5 % Glucose (40 %) in human serum albumin (20%)) and maintained at −150°C. Flow cytometry was used to identify and quantify lymphoma cells (GFP^+^) and CAR-T cells (GFP^-^, CD3^-^, CD4, CD8, CAR^+^). The list of antibodies used and dilutions used per staining are shown in supplement **Table S1** and gating strategies are shown in **Figure S2.** Analyses were performed using a Cytoflex S flow cytometer.

### Multiplex cytokine analysis

Cytokine concentrations from supernatants of cytotoxicity assays were determined using the LEGENDplex™ Human CD8/NK Panel Multi-Analyte Flow Assay (BioLegend), which allowed for the simultaneous detection of human Fas, FasL, IL-2, IL-4, IL-6, IL-10, IL-17A, Granulysin, Granzyme A, Granzyme B, Interferon (IFN)-γ, Perforin and Tumor necrosis factor (TNF)-α. The assay was performed according to the manufacturer’s instructions. Supernatants were diluted at 1:10 to 1:100 prior to the assay. Analysis was performed with the Data Analysis Software Suite for LEGENDplex™ version 2024-06-15 (BioLegend).

### Chromosomal aberrations and translocation analysis with CAST-Seq

Genomic DNA from double- or triple-edited primary T cells (day 14 after editing) was extracted using the DNeasy Kit (Qiagen, Hilden, Germany). Primer sequences used for CAST-Seq are listed in **supplement Table S2**. CAST-seq analyses were performed as previously described, with minor modifications [20–22]. In brief, 500 ng of genomic DNA was used as input material for each technical replicate. Libraries were prepared using the NEBNext Ultra II FS DNA Library Prep Kit for Illumina (New England Biolabs GmbH, Frankfurt, Germany). Enzymatic fragmentation of the genomic was aimed at an average length of 500–700 bp. CAST-Seq libraries were sequenced on a NovaSeq6000 using 150-bp paired-end sequencing (GENEWIZ/Azenta Life Sciences, Leipzig, Germany). For all CAST-Seq designs, two technical replicates were run and analyzed. All sites that were present and significant above background in at least one replicate are reported (**supplement Table S3**). For sites under investigation, the spacer sequence of the gRNA was aligned to the most covered regions for each site (±400 bp). Classification of off-target mediated translocations (OMTs) and the detection of gross ON-target aberrations were performed as previously described [22].

### High-throughput amplicon sequencing (HTS)

Amplicons of the TRAC/PDCD1/B2M target sites and the four putative off-target sites (OT1-4) identified by CAST-Seq were amplified using Q5® High-Fidelity DNA Polymerase (New England Biolabs GmbH, Frankfurt, Germany) and the primers listed in **supplement Table S2**. HTS libraries were generated using the NEBNext® Ultra™ II DNA Library Prep Kit for Illumina (New England Biolabs GmbH, Frankfurt, Germany) and NEBNext® Multiplex Oligos for Illumina® (New England Biolabs GmbH, Frankfurt, Germany) and sequenced on an Illumina NovaSeq 6000 by GENEWIZ/Azenta Life Sciences (Leipzig, Germany). The collected 2×150 bp paired-end reads were analyzed using CRISPResso2 [23]. The fraction of reads containing indels within the relevant editing window (±5 bp from the expected cut site) was reported.

### Statistical analyses

A t-test was used comparing the experimental groups with controls for data obtained in *in vitro* CAR-T cell killing assays and analyses of *in vivo* persistence of CAR-T cells and tumor development. The survival curves were compared using Log-rank (Mantel-Cox) test. The significance level was set to 0.05. Statistical analyses comparing three or more groups were carried out with one-way ANOVA using GraphPad Prism V10.5 software (GraphPad Software, La Jolla, CA).

## Results

### CRISPR/Cas9-generated TRAC^KI^-CD30CAR-T cells expand efficiently

The basic design of the HDRTs for integration of CAR constructs into the *TRAC* locus was described by Kath et al. [15] and adapted for incorporation of an improved signal peptide sequence [16]. The CAR constructs incorporate a CD28 costimulatory domain fused to a CD3ζ signaling domain (**Figure 1A**). To generate CD30-targeted CAR-T cells, we engineered HDRTs encoding mouse or human scFvs with varying heavy chain (VH) and light chain (VL) domain orientations. The CAR-T cell production initiated with gradient centrifugation separated PBMCs that were activated, electroporated, expanded, and analyzed by flow cytometry prior to cryopreservation (**Figure 1B**). For initial tests, we used the R-50×8 cuvettes for electroporation of eight different conditions (with or without RNPs or HDRTs). All HDRTs incorporating scFvs against CD30 (HRS-3, 5F11, 2H9) showed CAR expression by flow cytometric analysis for detection of the FcIg domain (**Figure 1C**). As expected, upon TRAC^KI^, expression of the TCR complex detectable via anti-CD3 staining was strongly reduced. Given that the anti-CD30 HRS-3 and 5F11 scFvs have been previously evaluated in clinical trials and showed an overall good potency and safety profile [7–9,12], CAR-T cells incorporating these scFvs were selected for further comparisons in triplicates. Compared to CD19CAR-T and HRS-3-CD30CAR-T cells, 5F11-CD30CAR-T cells demonstrated higher CAR expression at different time points after editing (Days 2, 7, and 9) (**Figure 1D**). The ratios of CD4^+^ and CD8^+^ *TRAC*^KI^-5F11-CD30CAR-T cells were comparable after *in vitro* expansion (**Figure 1E**). In summary, these results showed that CD30CAR-T cells expanded well and did not indicate fratricide effects (i.e., CAR-T cells eliminating putative CD30^+^ T cells after activation).

**Figure 1.**
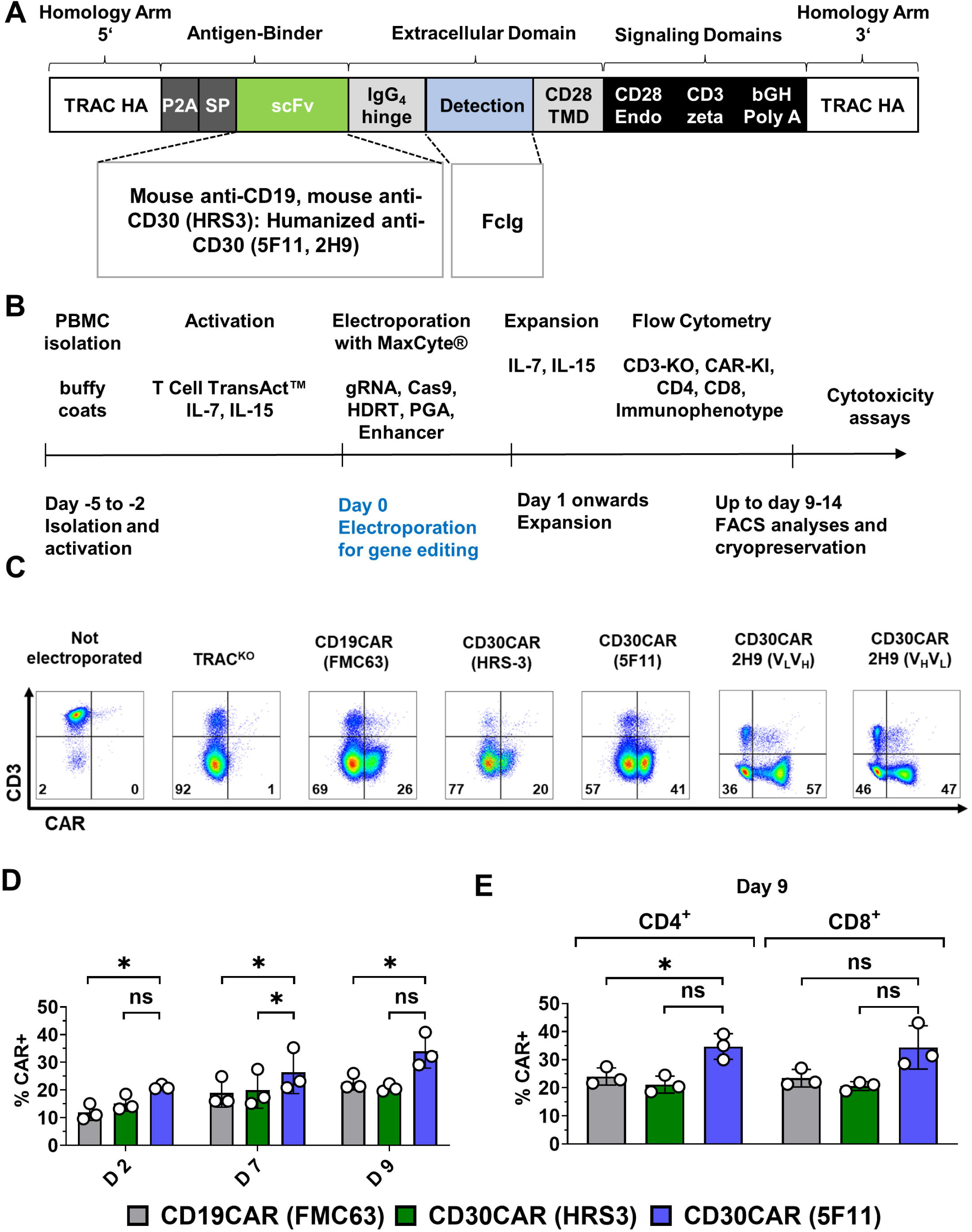
Generation and expansion of CD30CAR-T cells incorporating different single-chain antibody sequences (scFvs). (A) Scheme of the homology-directed repair template (HDRT) encoding the CAR and promoting specific integration into the *T cell receptor alpha constant (TRAC)* locus with *TRAC* homology arms (HA), transmembrane and endocytoplasmic CD28 domains, and CD3zeta signaling domain. The sequences for scFvs comprised a mouse anti-CD19, a mouse anti-CD30 (HRS-3) and human anti-CD30 sequences (5F11, 2H9). The detection domains included IgG4 hinge with IgG1-CH3 (FcIg). (B) Scheme of CAR-T cell production. After gradient separation, peripheral blood mononuclear cells (PBMCs) were activated for 2-5 days with anti-CD3 and anti-CD28 activating beads in the presence of IL-7 and IL-15. Prior to electroporation on day 0, the cells were mixed with gRNA, Cas9, HDRT, PGA, and an enhancer. After electroporation, cells were maintained in culture in the presence of IL-7 and IL-15 and harvested at different time points for analyses. (C) Flow cytometry analysis of CAR expression on 7 post electroporation for comparison of the different HDRTs/ scFvs, via detection of the CAR through the IgFc domain (representative data, n=1 donor). Not electroporated cells and cells electroporated with RNPs but no HDRT (*TRAC*^KO^) are shown as controls. Lack of CD3 detection indicates TCR knockout. (D) Flow cytometry analysis of CAR expression on days 2, 7, and 9 post electroporation as quantified data comparing the expression of the CARs incorporating different scFvs (CD19CAR in grey, HRS-3-CD30CAR in green, and 5F11-CD30CAR in blue) and detected via the FcIg domain (n=3 donors). (E) Flow cytometry analysis of CAR expression on day 9 post electroporation separated for CD4^+^ and CD8^+^ cells as quantified data comparing the expression of the CARs incorporating different scFvs (CD19CAR in grey, HRS-3-CD30CAR in green, and 5F11-CD30CAR in blue) (n=3 donors). Statistical significance is annotated: n.s.: not significant, *: p>0.05.

### 5F11-CD30CAR-T cells efficiently kill EBV-infected lymphoma cells *in vitro*

We have previously demonstrated the potency of gene-edited CD19CAR-T cells to kill EBV-infected CD19^+^ Burkitt lymphoma cells [16]. For the experiments described here and further below, we evaluated if CD30CAR-T cells could kill CD30^+^ cell lines derived from HL patients (L591 and L540), from TCL patients (Myla), or from primary B cells infected with EBV and transformed into LCLs recapitulating EBV-LPD. L591 (EBV positive), L540 (EBV negative), and Myla (EBV negative) cell lines were transduced with a lentiviral vector co-expressing firefly luciferase (fluc, for bioluminescence detection) and green fluorescent protein (GFP, for flow cytometry) (**Figure 2A, supplement Figure S1**). LCLs were infected with either the latent EBV strain B95.8 or the lytic EBV strain M81. Quantitative PCR analysis was used to confirm detectable EBV viral load (**Figure 2B**). Cytotoxicity assays demonstrated robust anti-tumor activity of CD30-targeted CAR-T cells derived from three independent donors against L591 cells, with lysis ranging from 60% to 100% at effector-to-target (E:T) ratios of 3:1. Notably, cytolytic capacity exhibited donor-dependent variability, with donor 1 achieving complete target cell lysis (100%) at 1:1 E:T ratio (**Figure 2C**). Comparable cytotoxic potentials were observed when CAR-T cells generated from donor 1 were co-cultured with LCLs infected with either EBV strain B95.8 or M81 (**Figure 2D**). We decided to proceed with 5F11-CD30CAR-T cells as the CAR expression and expansion were superior to HRS-3-CD30CAR-T cells.

**Figure 2:**
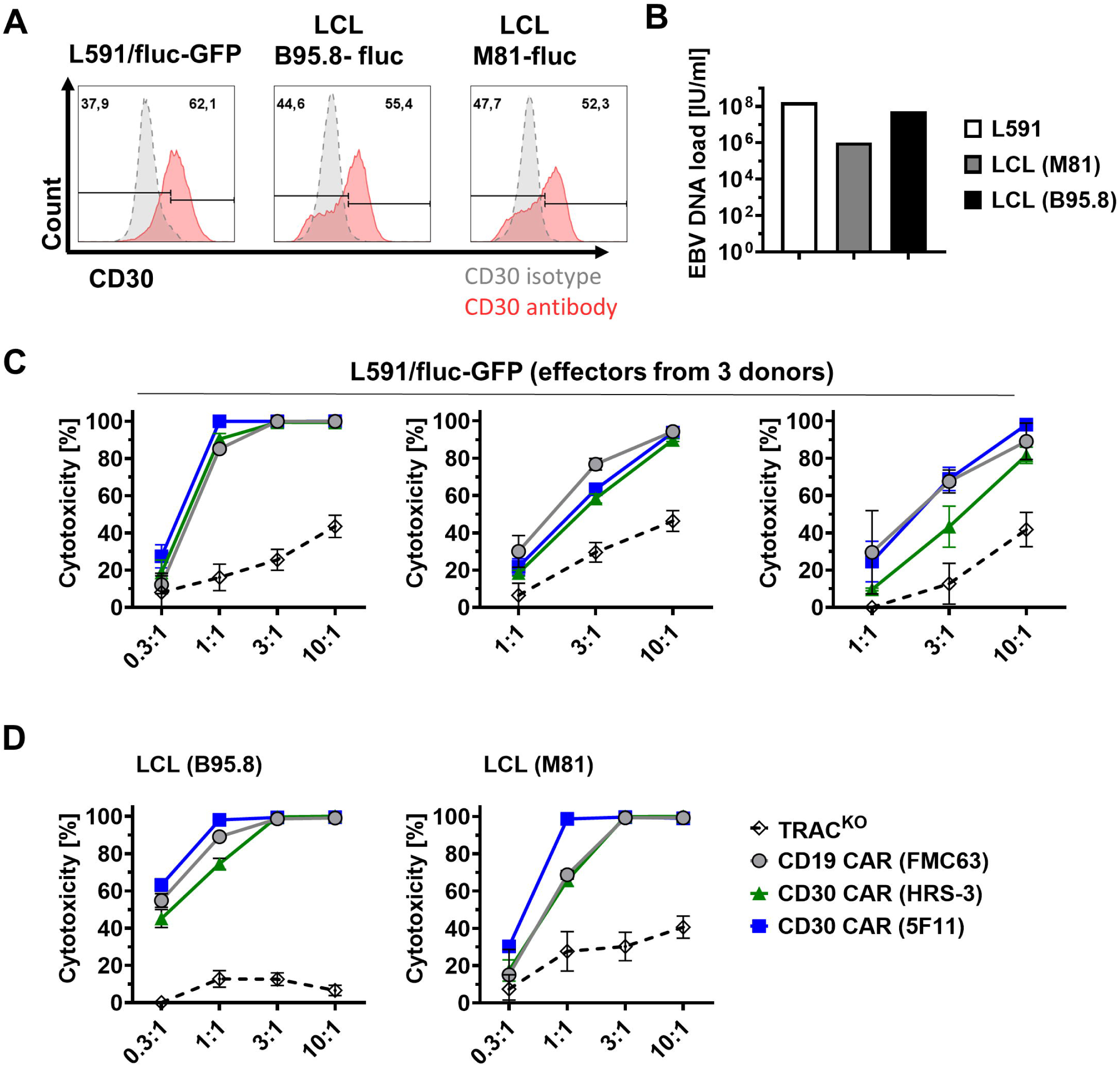
Killing potency of CD30CAR-T cells against EBV-infected cells. (A) Detection of CD30 on EBV^+^ cells. Histograms show staining for flow cytometry analyses with anti-CD30 antibody (red) versus isotype (grey). (B) Analysis of EBV load in L591/fluc-GFP cell line and lymphoblastoid cell lines (LCL)/fluc cell lines via quantitative PCR (infectious units (IU) per milliliter (ml)). (C) Bioluminescence-based cytotoxic assay to determine potency. CAR-T cells were cultured for 72 h with L591 cells expressing fluc-GFP at different Effector:Target (E:T) ratios (0.3:1, 1:1, 3:1, 10:1) (n=3 donors). (D) CAR-T cells were cultured for 72 h with LCLs expressing fluc at different E:T ratios (0.3:1, 1:1, 3:1, 10:1) (n=1 donor). Cytotoxicity was calculated based on remaining bioluminescence detectable after co-culture. TRAC^KO^ hatched line, CD19CAR in grey, HRS-3-CD30CAR in green, and 5F11-CD30CAR in blue.

### Incorporation of a tCD34 spacer: Improved CD30CAR detection, accumulation of expanded naïve and central memory cells and potent killing functions

Detection of the CARs by flow cytometry analyses via the FcIg spacer showed limitations. The conjugated polyclonal antibody provided low fluorescence signals, and TRAC^KO^ and CAR^KI^ cell populations could not be well distinguished (**Figure 1C**). As an optimization of the 5F11-CD30CAR construct, we explored a previously described tCD34 spacer/detection domain [19] with some sequence and structural modifications (**Figure 3A)**. Flow cytometric analyses performed with a single fluorochrome-labelled monoclonal antibody revealed a clear separation of the TRAC^KO^ and CAR^KI^ populations (**Figure 3B**). For 3 donors analyzed, expanded CD4^+^ and CD8^+^ CD30CAR⁺ T cells showed an average of 50% CAR-positivity and exhibited up to a tenfold expansion from day 5 to 12 after electroporation (**Figure 3C**). Immunophenotypic analyses indicated that most expanded CD8⁺ CD30CAR-T cells were classified as naïve (N), while most expanded CD4⁺ CD30CAR-T cells were classified as central memory (CM) (**Figure 3D).** On day 5 after electroporation, CD8⁺ CAR-T cells were mostly CM, and expansion until day 12 somehow shifted them to an N phenotype. On the other hand, CD4⁺ CD30CAR-T cells displayed a stable CM phenotype from days 5 to 12 after electroporation (**Figure 3E**). Functional potency assays demonstrated > 90% cytotoxic activities of CD30CAR-T cells against the L540 cell line at an E:T ratio of 1:1 (3 donors). For the EBV-positive L591 cell line, > 90% cytotoxic activity of CD30CAR-T cells was observed at E:T ratios of 3:1 (**Figure 3F**). Therefore, 5F11-CD30CAR-T with the tCD34 detection domain was selected for further *in vivo* studies.

**Figure 3:**
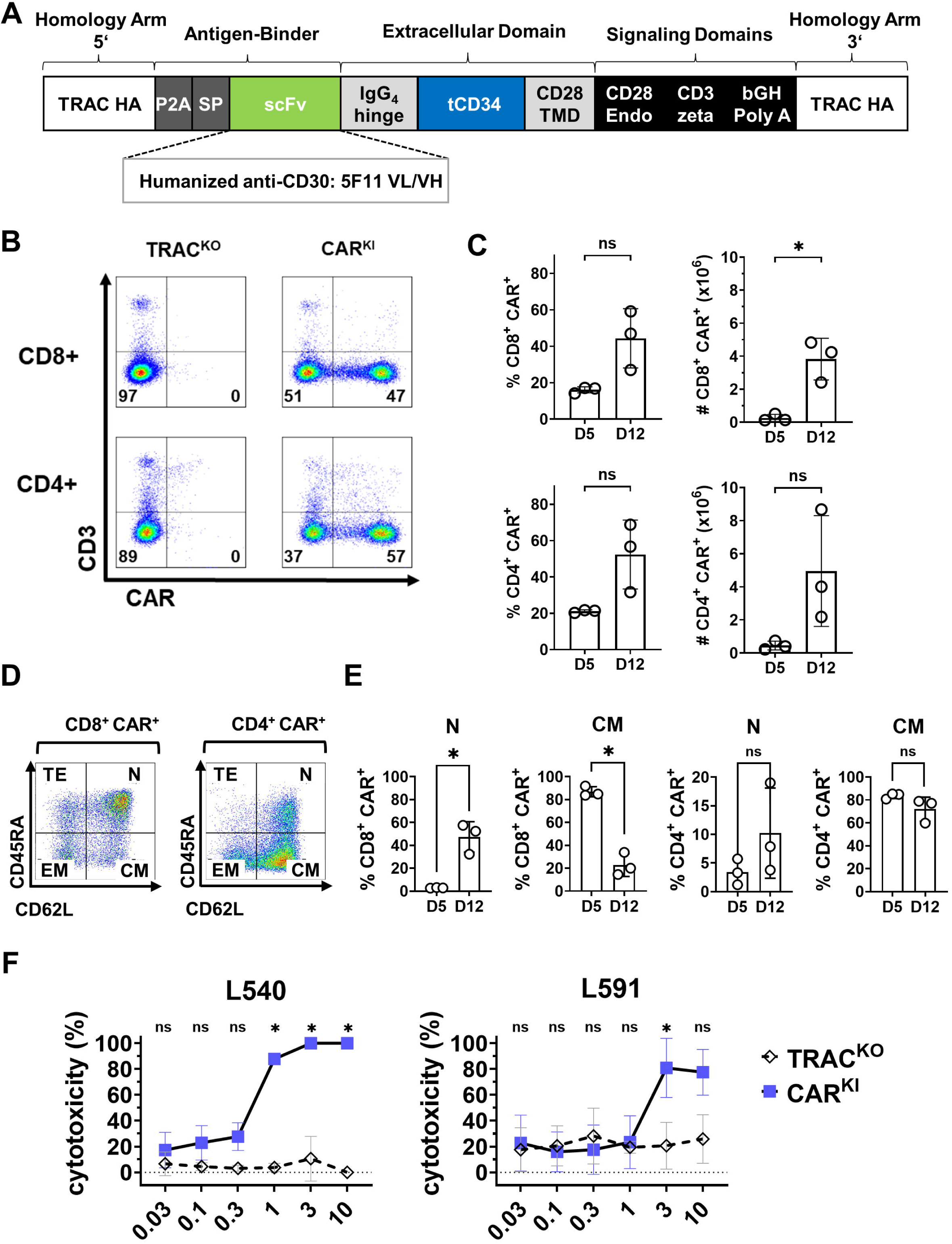
Improvement of CAR expression and detection with the truncated CD34 detection domain (tCD34). (A) Schematic overview of HDRT incorporating the tCD34 hinge for 5F11-CD30CAR-T cell generation. (B) Analysis of CAR expression. Representative flow cytometry analyses of CAR expression on day 12 after electroporation. TRAC^KO^ cells, electroporated RNPs but no HDRT, are shown as controls. Lack of CD3 detection indicates TCR knock-out. (C) Percentages and absolute numbers of CD4^+^ and CD8^+^ CD30CAR-T cells detectable on day 5 and day 12 after electroporation showing expansion (n=3 donors). (D) Immunophenotypes of CD30CAR-T cells defined by expression of CD62L and CD45RA. Representative flow cytometric analysis of phenotypes of CD8^+^ and CD4^+^ CAR-T cells on day 12 after electroporation. N: Naïve, CM: central memory, EM: effector memory, TE: Terminal effector. (E) Quantified frequencies of naïve (N) and central memory (CM) phenotypes within CD8^+^ and CD4^+^ CD30CAR-T cells on day 5 and 12 after electroporation indicating their accumulation upon culture (n=3 donors). (F) Effectors (TRAC^KO^ hatched lines and CAR^KI^ blue lines) were cultured for 72 h with targets L540/fluc-GFP (EBV-negative) and L591/fluc-GFP (EBV-positive). Assay was performed with different Effector : Target ratios (0.03:1, 0.01:1, 0.3:1, 1:1, 3:1, 10:1) (n=3 donors). Cytotoxicity was calculated based on bio-luminescence remaining in target cells. Statistical significance is annotated: n.s.: not significant, *: p>0.05.

### TRAC^KI^5F11-CD30CAR-T cells showed *in vivo* therapeutic responses in the L540 xenograft mouse model

Production of CD30CAR-T cells was upscaled using the R-1000 cuvettes for electroporation in order to obtain enough cells to evaluate the functionality of CD30CAR-T cells *in vivo*. The expanded cell product commonly exhibited CD30CAR^+^ T cell frequencies of >70% (**Figure 4A**) and the cells were cryopreserved. It is known that HL is a complex tumor entity highly interacting with the tumor microenvironment (TME). Due to the lack of a *bonafide* xenograft model, we employed L540 cells that engraft well in immunodeficient mice and were previously used to test the *in vivo* therapeutic efficacy of the 5F11 monoclonal antibody [24]. We adapted the model using immunodeficient NSG-DKO mice, which lack the expression of mouse MHC class I and class II molecules and therefore mount less xenograft reactivity with infused human T cells [25]. L540/fluc-GFP cells were injected i.v., and tumor engraftment was confirmed by BLI 4 days post-injection for all mice. On the next day, 1×10⁶ CD30CAR-T cells were administered i.v. (**Figure 4B**). Monitoring of the mice challenged with L540 tumor cells (treated with PBS) showed engraftment on day 4 and afterwards a rapid tumor progression from the anatomical regions of liver and legs to all body, indicated by a high bioluminescence signal. From week 4 onwards, all mice started showing rapid weight loss (**Figure 4D upper panel**) and signs of paralyzed legs. All mice receiving tumor only had to be terminated within 8 weeks after challenge as they reached the unfit score (**Figures 4C, 4D upper panel**). Accordingly, mice challenged with tumor showed continuous and rapid increase in the quantified bioluminescence signals (**Figures 4C, 4E upper panel**). In contrast, most mice treated with CD30CAR-T cells showed weight loss at later timepoints (6-10 weeks after tumor challenge) (**Figure 4D**) and a continuous decrease of bioluminescence signals (**Figures 4C, 4E lower panel**). Some CD30CAR-T-treated mice showed bioluminescence signals persisting in the anatomic region of the brain (**Figure 4C**) and paralysis, which might indicate tumor infiltration in the central nervous system. Although all CD30CAR-T-treated mice succumbed, the cell therapy treatment showed significant effects in prolonging survival (**Figure 4F**). In sum, CD30CAR-T cells showed *in vivo* therapeutic potency in the L540 xenograft model.

**Figure 4:**
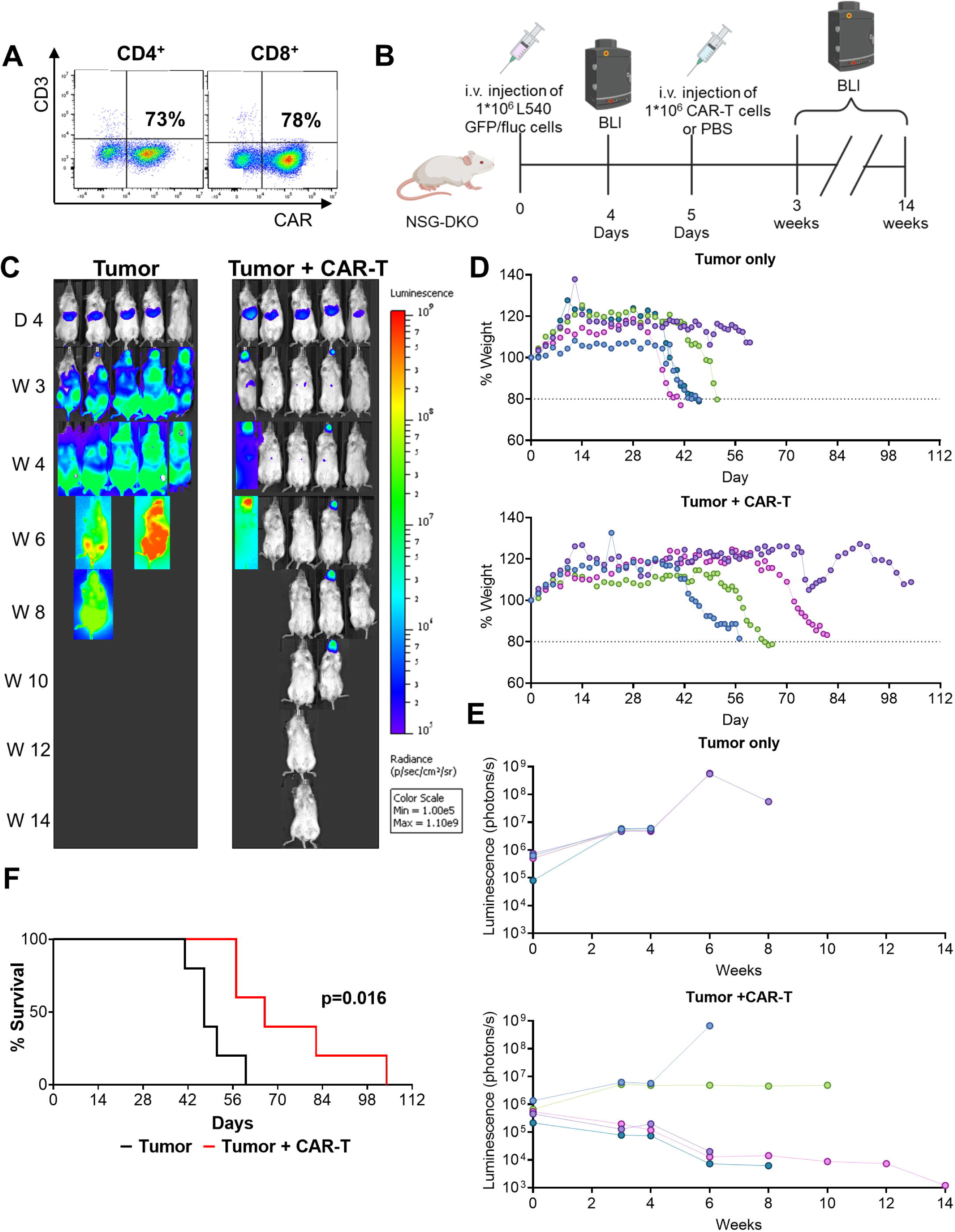
Evaluation of TRAC^KI^-5F11-CD30CAR T efficacy *in vivo*. (A) Flow cytometry analysis of CAR-T cells used in the experiment. The plots display high (>70%) CAR expressing cells within CD4^+^ and CD8^+^ populations. (B) Scheme of experiment: NSG-DKO mice were injected i.v. with 1×10^6^ L540 GFP/fluc cells. Bio-luminescence imaging (BLI) analyses were performed on day 4 after tumor challenge to confirm the baseline tumor engraftment. On day 5, five mice were injected i.v. with 1×10^6^ CD30CAR-T cells, and five mice received PBS. The mice were monitored for weight and the tumor progression bi-weekly via BLI. (C) BLI pictures obtained on day 4 after tumor implantation and afterwards bi-weekly. One treated mouse survived for more than 100 days after tumor challenge. High BLI signals were observed systemically in PBS-treated mice, whereas the mice that received CAR-T cells showed sporadic weak signals in the anatomical region of the brain. (D) Monitoring of weight changes over time showing rapid weight loss in mice challenges with tumor and treated with PBS and slower weight loss in mice challenged with tumor and treated with CD30CAR-T mice. (E) Sequential BLI measurements quantified as photons/s/cm^2^/sm. Mice treated with CD30CAR-T cells show a major reduction in luminescence signal compared with mice treated with PBS. (F) Kaplan-Meier survival curves. Mice treated with CD30CAR-T cells displayed significantly longer survival. Statistical significance is annotated: n.s.: not significant, *: p>0.05.

### L540 tumors homed in bone marrow, and CD30CAR-T cells showed persistence in bone marrow and spleen with high CD4/CD8 ratios

Terminal analyses of different tissues by flow cytometry for detection of L540/fluc-GFP showed homing of tumor cells in bone marrow, with a significantly reduced tumor burden in mice treated with CD30CAR-T cells (**Figure 5A, for gating, see supplement Figure S2)**. To characterize the homing and persistence of CD30CAR-T cells in the mice, blood and tissues were further analyzed by flow cytometry. CD30CAR-T cells were detectable in blood samples up to 8 weeks after tumor challenge (**Figure 5B**). Additionally, the bone marrow of treated mice showed persistent CD30CAR-T cells, with a significant preponderance of CD4^+^ cells over CD8^+^ T cells (ca. 60 /40 %) (**Figure 5C**). A similar trend was observed for CD30CAR-T cells homing in the spleen, although the values did not reach statistical significance (**Figure 5D**). In summary, lower tumor burden upon CD30CAR-T cell therapy was associated with detectable CAR-T cells in blood and bone marrow, and CD4^+^ cells showed higher persistency.

**Figure 5:**
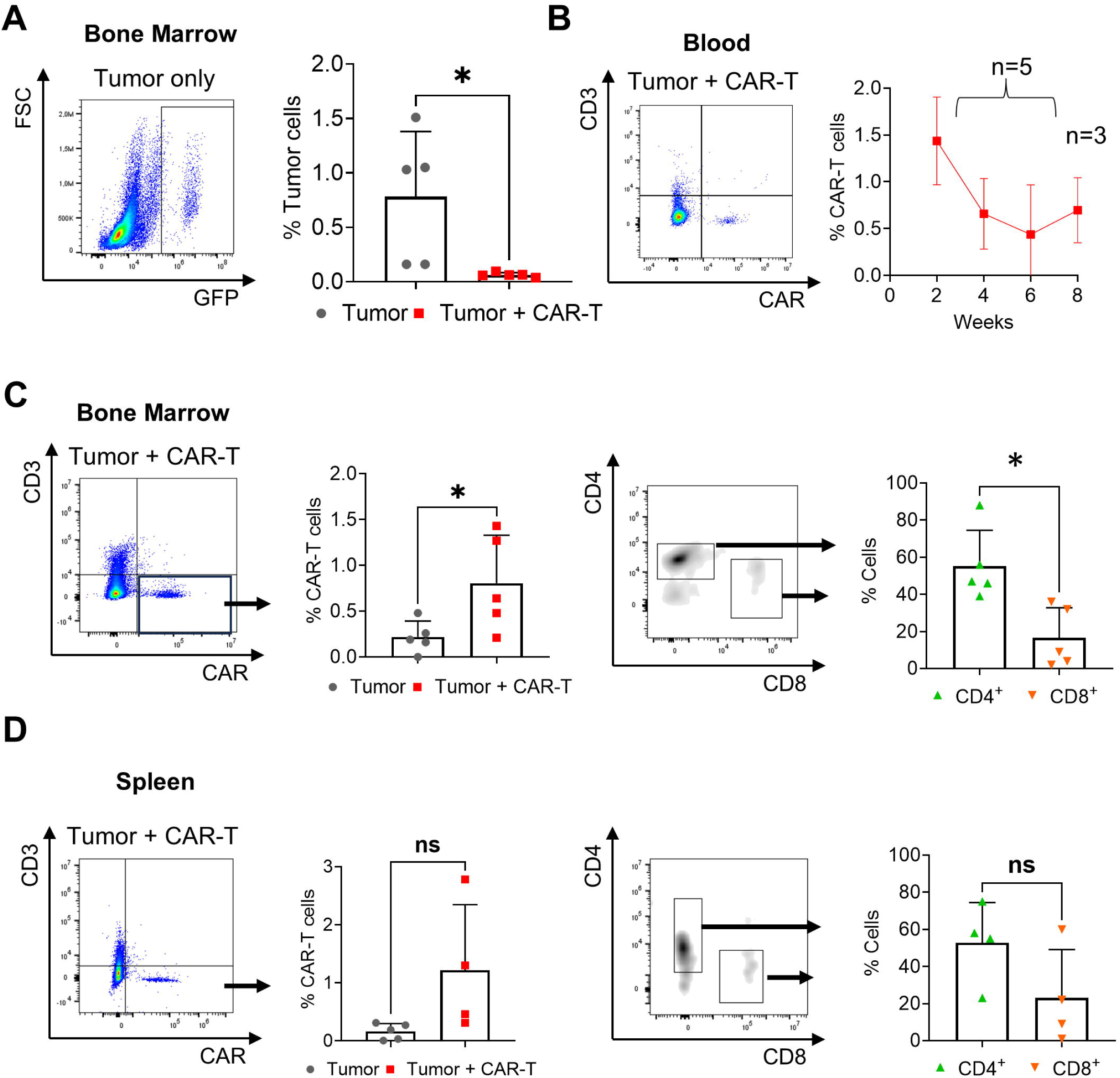
Tissue analyses associating tumor reduction with high persistency of TRAC^KI^-5F11-CD30CAR-T cells. (A) Flow cytometry analyses of bone marrow showing infiltration of GFP^+^ tumor cells (representative plot on the right side) and significant tumor reduction in CD30CAR-T treated mice and analyzed at terminal endpoint (n=5 for each cohort, tumor only: grey dots; tumor + CAR-T: red dots). (B) Flow cytometry analyses of peripheral blood showing presence of CAR-T cells (representative plot on the right side) Analyses of blood samples collected after tumor challenge until week 6 (n=5) or week 8 (n=3). (C, D) Flow cytometry analyses of bone marrow (C) or spleen (D) showing presence of CAR-T cells (representative plots on the right side and plots in the middle to differentiate CD4^+^ and CD8^+^cells) Tissues were collected at terminal endpoint showing detectable CD30CAR-T cells (left panels) and that CD4^+^ cells showed higher persistency than CD8^+^ cells (tumor only: n=5; tumor + CAR-T: n=5). Statistical significance is annotated: n.s.: not significant and *: p>0.05.

### TRAC^KI^CD30CAR-PD-1^KO^-T cells are feasible and functional

Lymphoma cells commonly express the programmed death ligand 1 (PD-L1), which is known to interact with PD-1 on T cells and cause their inhibition. L540, Myla, and L591 cell lines used in this study expressed PD-L1 (**supplement Figure S4**). As a possible strategy to improve the potency of CD30CAR-T cells, we evaluated if the KO of the *PDCD1* gene (that encodes the immune checkpoint PD-1) could protect CAR-T cells from exhaustion. Four different guide RNAs targeting *PDCD1* were designed and evaluated for CRISPR/Cas KOs using activated PBMCs from two donors. The gRNA 1 was selected, showing the highest knockout efficiency of PD-1 expression 12 days post-electroporation of PBMCs (**supplement Figure S5**). The PD-1^KO^ gRNA was therefore added to the RNP complex used to generate CD30CAR-T cells as an all-in-one KI/KO gene editing approach. Analyses of the CD8^+^ and CD4^+^ cells on day 12 after gene editing showed no detrimental effects of including the additional PD-1^KO^ gRNA on the generation of CD30CAR-T cells (**Figure 6A**). Knockout of PD-1 expression was confirmed, showing a more apparent effect on CD4^+^ CD30CAR-T cells (**Figure 6B**). PD-1^KO^ did not affect the *in vitro* expansion of CD8^+^ or CD4^+^ CD30CAR-T cells between days 5 and 12 post-editing (**Figure 6C**). In addition, PD-1^KO^ did not alter the increase in N immunophenotypic characteristics of CD8^+^ CD30CAR-T cells and stable CM immunophenotypic characteristics of CD4^+^ CD30CAR-T cells (analyses on days 5 and 12 post-editing) (**Figure 6D**). Therefore, the all-in-one PD-1^KO^ editing approach was feasible and did not change the expansion or overall immunophenotypic characteristics of CD30CAR-T cells.

**Figure 6:**
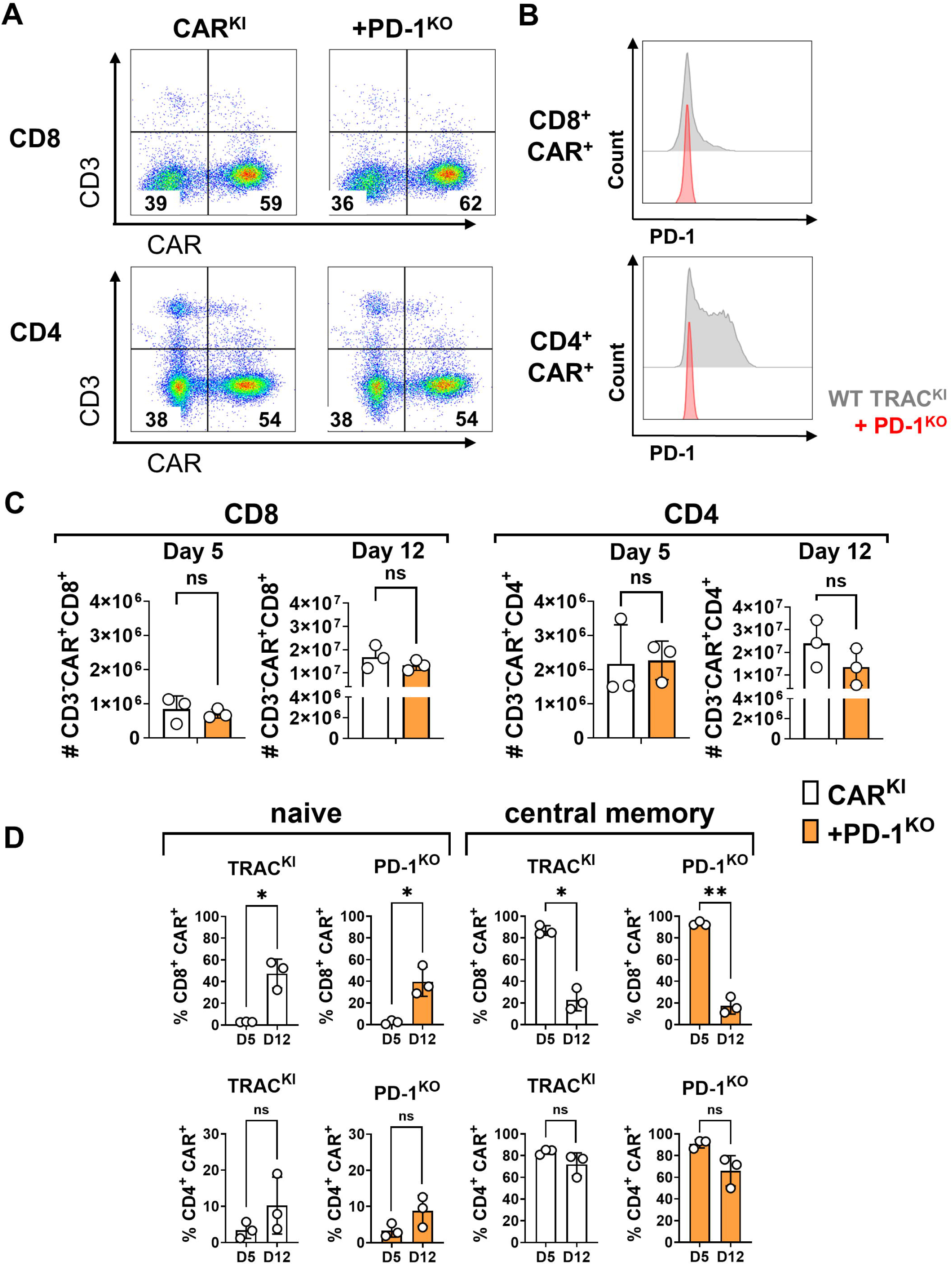
All-in-one PD-1^KO^ and TRAC^KI^-CD30CAR-T generation. (A) Flow cytometry analyses of CAR expression within CAR^KI^-T and CAR^KI^ PD-1^K^-T cells on day 12 after electroporation. Representative flow cytometry plots showing CD8^+^ and CD4^+^ CD30CAR-T cells. (B) CD8^+^ and CD4^+^ CD30CAR-T cells co-electroporated with gRNA 1 showed efficient PD-1^KO^. Representative flow cytometry analyses. (C) Quantitative analyses showing expansion of TRAC^KI^ or additional PD-1^KO^ CD30CAR-T cells separated between CD8^+^ and CD4^+^ populations at days 5 and 12 after electroporation (n=3 donors). (D) Quantitative analyses showing naïve or central memory phenotypes for TRAC^KI^ or additional PD-1^KO^ CD30CAR-T cells separated between CD8^+^ and CD4^+^ populations from day 5 to 12 after electroporation (n=3 donors). Statistical significance is annotated: n.s.: not significant, *: p>0.05, **: p>0.005.

### TRAC^KI^CD30CAR combined with PD-1 and β2M knock-outs is feasible, but affects the cell expansion

Heavily treated lymphoma patients sometimes develop lymphopenia, and therefore generation of autologous CAR-T cells can become a challenge. Thus, KO of the gene expressing the β2M molecule in CAR-T cells could potentially facilitate the procurement of third-party allogeneic donors, because loss of HLA-I would lower immune rejection by allo-reactive CD8^+^ T cells. Therefore, we evaluated the feasibility of the simultaneous HLA-I^KO^ and PD-1^KO^ via gene editing. HLA-I^KO^ did not impair generation or expansion of CD4^+^ and CD8^+^ knock-in CAR-T cells (**Figure 7A-C**). Flow cytometric analyses confirmed complete KO of HLA-I and PD-1 expression within the CD4^+^ and CD8^+^ fractions of CAR-T cells (**Figure 7B)**. A trend of reduced expansion of the CAR-T cells following simultaneous HLA-I^KO^ and PD-1^KO^ was observed (**Figure 7C**). Taken together, these data indicated that HLA-I^KO^ alone was feasible, but combined with PD-1^KO^, affected the cell expansion.

**Figure 7:**
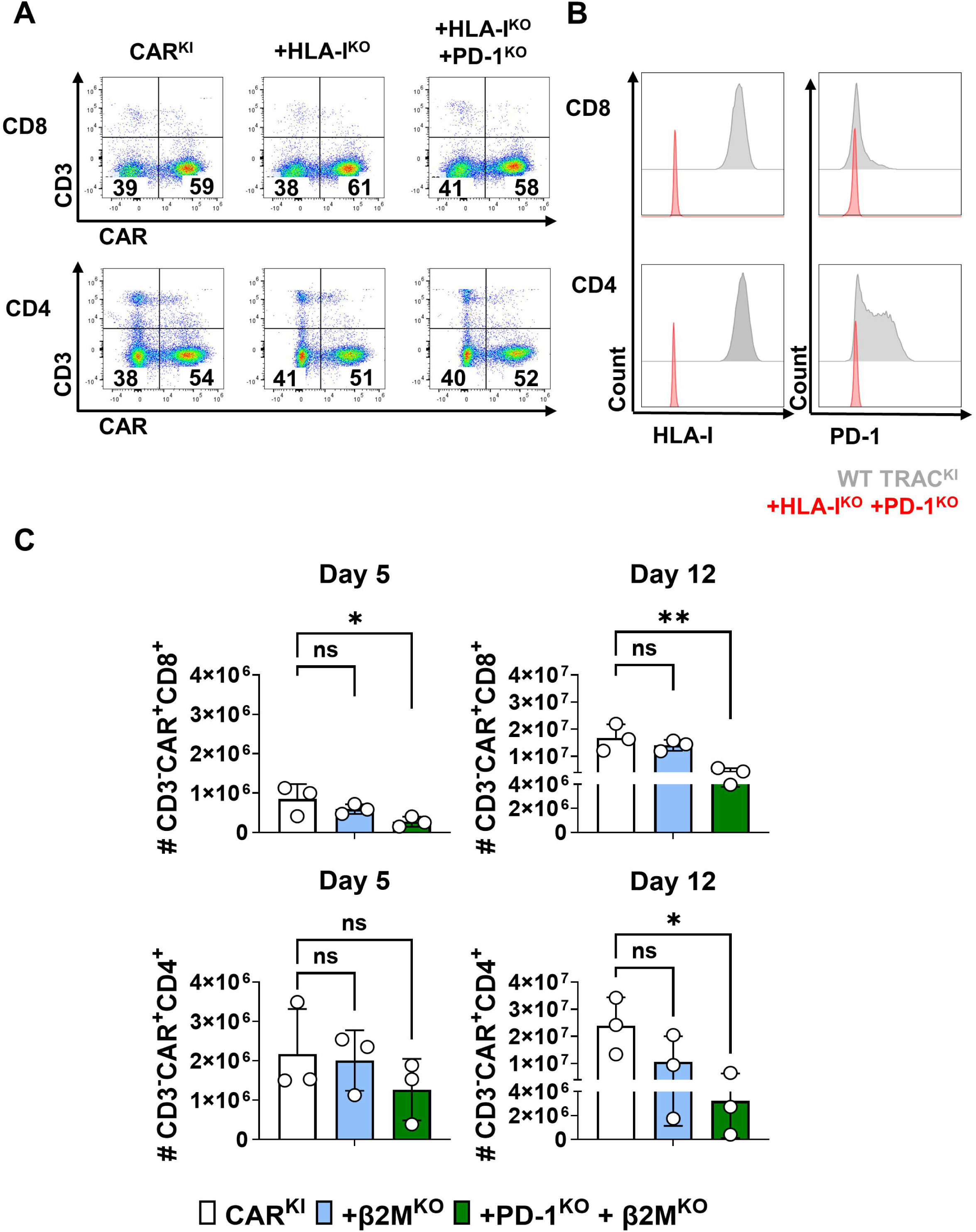
All-in-one PD-1^KO^ and HLA-I^KO^ and TRAC^KI^CD30CAR-T generation is feasible, but affects cell expansion. (A) Representative flow cytometry analyses of CAR^KI^, CAR^KI^ HLA-I^KO^ and CAR^KI^ HLA-I^KO^ PD-1^KO^ T cells on day 12 after electroporation, showing CD8^+^ and CD4^+^ CD30CAR-T cells. (B) CD8^+^ and CD4^+^ CD30CAR-T cells co-electroporated with gRNAs for simultaneous HLA-I^KO^ and PD-1^KO^. Representative flow cytometry analyses. (C) Quantitative analyses showing expansion of CD8^+^ and CD4^+^ CD30CAR-T cells with CAR^KI^, CAR^KI^ HLA-I^KO^ or CAR^KI^ HLA-I^KO^ PD-1^KO^ at days 5 (left panels) and 12 8right panels) after electroporation (n=3 donors). Statistical significance is annotated: n.s.: not significant, *: p>0.05, **: p>0.005.

### CD30CAR-T cells with HLA-I^KO^ and PD-1^KO^ show *in vitro* potency against different lymphomas

Cytotoxicity assays were performed to evaluate the functional effects of the different KOs using L540, L591, and Myla lymphoma target cell lines expressing PD-L1 (**supplement Figure S4**). Cytotoxicities of CD30CAR-T cells generated from three donors were highly variable among the donors (**Figure 8A**). Overall, knockout of β2M or PD-1 expression did not substantially improve or inhibit the cytotoxicity potential of CD30CAR-T cells, although subtle differences depending on the target cells and editing modalities could be observed (**Figure 8A**). The benefit of PD-1 disruption in CD30CAR-T cells to improve the cytotoxicity was mostly marginal and observed at lower E:T ratios. For L540 and L591 target cells exposed to CD30CAR-T cells at 1:1 E:T ratios, a slightly higher trend for an increased cytotoxicity of CAR-T cells with PD1^KO^ was observed (**Figure 8B)**. These results may reflect the fact that CD8^+^ CD30CAR-T cells showed a low expression of PD-1 (**Figures 6B, 7B**) and its KO did not consistently improve their cytotoxic T lymphocyte (CTL) function.

**Figure 8:**
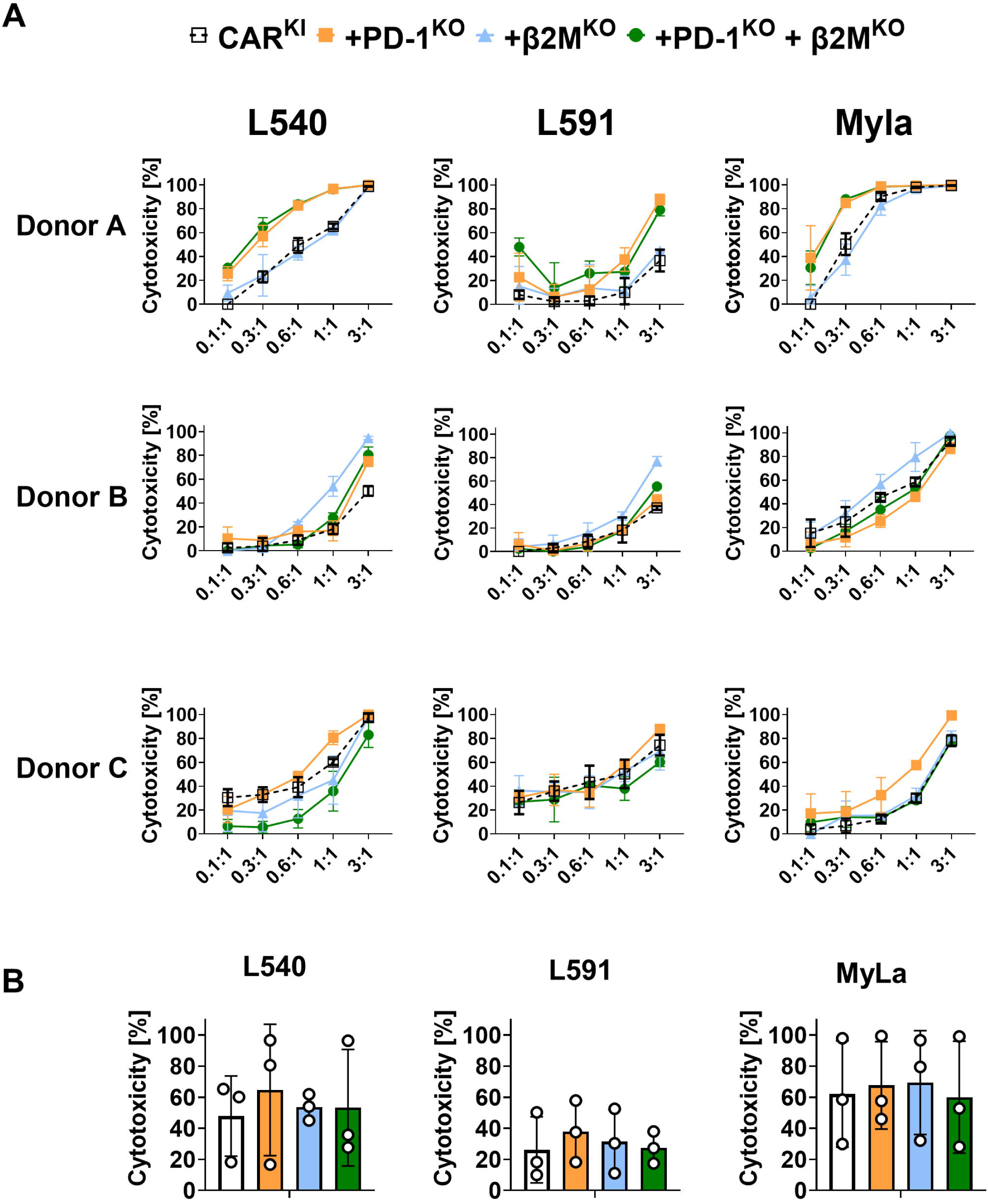
CD30CAR-T cells with PD1^KO^, HLA-I^KO^ and HLA-I^KO^ PD-1^KO^ retain functionality, but do not show improved cytotoxicity. (A) Effector cells were generated with 3 donors (A, B, C), and were expanded for 12 days after electroporation. The cells were cryopreserved, thawed and then co-cultured with L540, L591 or Myla cells co-expressing fluc-GFP. Effector : Target (E:T) ratios 3:1, 1:1, 0.6:1, 0.3:1 and 0.1:1 were set up and cytotoxicity was detected by bio-luminescence analyses after 72 h of co-culture. (B) Cytotoxicity at an Effector : Target ratio of 1:1. Data were merged from 3 different donors. Statistical analyses showed no significant differences among arms.

### CD30CAR-T cells show elevated secretion of proinflammatory cytokines upon PD-1^KO^

Since CD4^+^ CD30CAR-T cells showed a high expression of PD-1 (**Figures 6B, 7B**), we next evaluated if proteins secreted in the cell culture medium after exposure to the target cells could indicate effects on T helper (Th) function. To characterize the protein secretion profiles of the multiply edited CAR-T cells, culture supernatants of cultures performed at 1:1 E:T ratios were analyzed by immunodetection using bead assays. The proteins Granzyme A, Granzyme B, interferon-γ (IFN-γ), and Fas ligand (FasL) were detectable at variable levels and higher concentrations in supernatants of CD30CAR-PD-1^KO^-T cells compared with CD30CAR-T cells (**Figure 9**). Overall, the magnitude of protein secretion effects was higher for the L540 and L591 co-cultures than for Myla. The concentration of these proteins when comparing co-cultures of CD30CAR-HLA-1^KO^-T cells with CD30CAR-T cells remained mostly unchanged. The combined CD30CAR-PD-1^KO^HLA-1^KO^-T cells showed an intermediate pattern between CD30CAR-PD-1^KO^-T cells and CD30CAR-HLA-1^KO^-T cells. In conclusion, PD-1^KO^ in CD30CAR-T increased the secretion of some proinflammatory proteins when the cells were exposed to PD-L1-positive targets.

**Figure 9:**
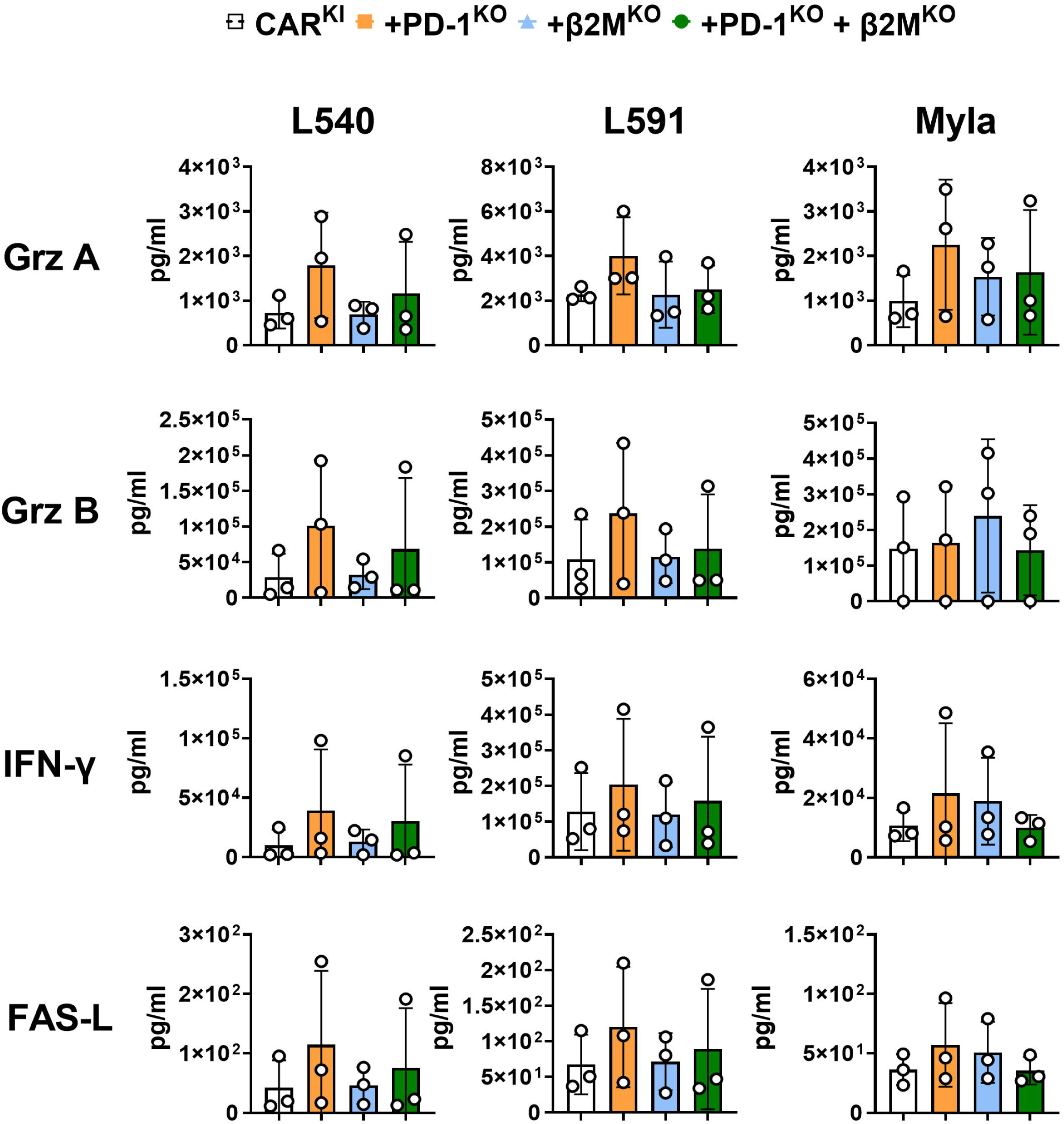
CD30CAR-T cells show elevated secretion of proinflammatory cytokines upon PD-1^KO^. Cytokine analysis of Granzyme A (Grz A), Granzyme B (Grz B), Interferon-γ (IFN-γ), and Fas-ligand (FAS-L) in supernatants from cytotoxic assays. Cytotoxic assays were performed for 72h coculturing CD30CAR-T cells (white) and CD30CAR-T cells with additional knock outs of PD-1 (orange), β2M (blue) and PD-1 and β2M (green) against L540, L591 or Myla target cell lines at an Effector to Target (E:T) ratio of 1:1. Supernatants were taken and frozen at −80°C until analysis. CAR-T cells were produced in 3 biological replicates. Statistical analyses showed no significant differences among arms.

### CD30CAR-T cells simultaneously edited at *TRAC*, *PDCD1,* and *β2M* loci show chromosomal translocations

To evaluate possible genomic instability, including chromosomal rearrangements and large on-target aberrations, we performed CAST-Seq on gene-edited CAR-T cells with double or triple KOs in TRAC, PDCD1, and/or β2M in combination with targeted KI of the CD30CAR on day 14 after editing. Chromosomal translocations were detected between the on-target sites (TRAC/PDCD1/β2M) two weeks post editing. Moreover, a total of four off-target (OT) activity-mediated translocations with sites in intronic regions were detected in ≥2 replicates (**Figure 10A**; **supplement Table S3**). Two of the four putative OT sites – OT1 on chromosome 15 (with sequence homology to the PDCD1-targeted sgRNA) and OT2 on chromosome 10 (homology to the β2M sgRNA) were found across samples (**Figure 10B**). OT1 and OT2 diverged from the sgRNA sequences by a single DNA bulge or two DNA bulges and a single mismatch in the PAM-distal region, respectively. To assess the extent of on-target and OT activities, all seven sites were subjected to NGS. Mean indel frequencies at on-target sites reached 91% for TRAC, 99% for PDCD1, and 99% for β2M (**Figure 10C**). We further confirmed off-target activity at OT1 (35% alleles with indels) in all samples that included the PDCD1-targeted RNP, as well as OT2 in all samples edited with the β2M-targeted Cas9 complex (1.0% indels). Indel formation at OT3 and OT4 was not above background. As expected, large aberrations at both examined on-target sites were detected (**supplement Figure S6**). Taken together, triple editing induced stable chromosomal aberrations, the biological impact of which must be further evaluated.

**Figure 10:**
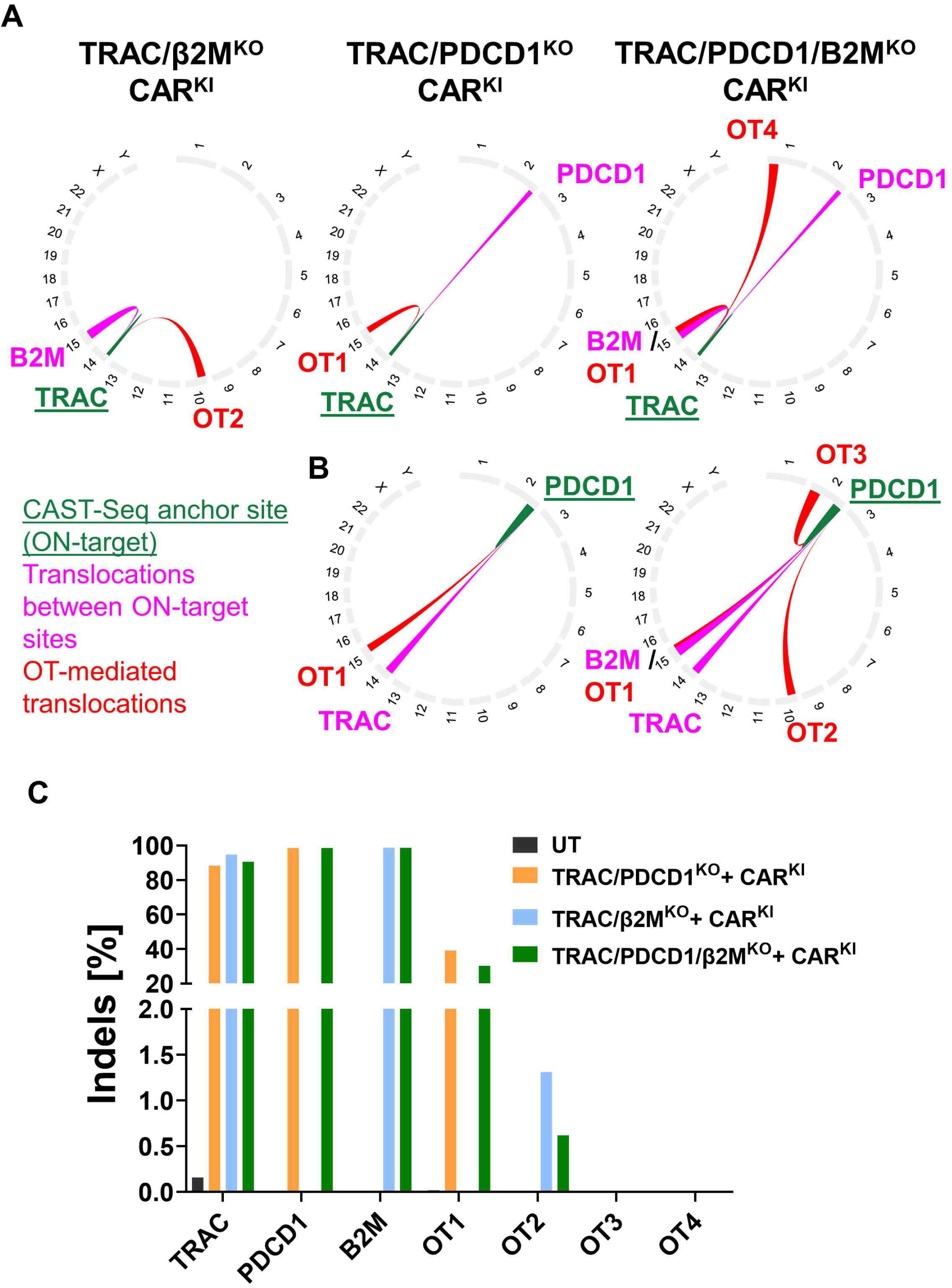
Chromosomal rearrangements in multiplex edited CD30CAR-T cells. Chromosomal rearrangements were evaluated by CAST-Seq on day 14 after electroporation employing at least two CAST-Seq libraries. Shown are chromosomal translocations with (A) *TRAC* and (B) *PDCD1* target sites. Green line, ON-target aberrations; magenta, chromosomal translocations between target sites; red lines, off-target activity-mediated translocations. (C) Indel frequencies as determined by targeted amplicon sequencing of ON-target and OFF-target sites nominated by CAST-Seq.

## Discussion

Current standard of care for cHL results in excellent outcomes in youngers and unsatisfying results in elderly, with a considerable risk of morbidity due to long-term side effects from frontline chemotherapies. Thus, new less toxic approaches such as immune and cell therapies still remain relevant. PD-1 blockade combined with brentuximab vedotin, an anti-CD30 antibody-drug conjugate (ADC), has shown impressive results in multiply relapsed cHL patients. Nevertheless, newer cellular immunotherapy approaches are also warranted due to the immunologically dysfunctional cHL tumor milieu, revealing a regulatory T-cell–rich and exhausted T-effector microenvironment [26]. CD30 expression is also often observed in virus-infected cells such as human T-cell leukemia virus type 1 (HTLV-1, resulting in highly refractory adult T-cell leukemia) and EBV (associated with aggressive types of B-cell lymphomas) [2,27]. CD30 is universally expressed in anaplastic large cell lymphoma (ALCL) and up to 30% of other peripheral T cell lymphomas (PTCLs) [28]. Despite the proven clinical success of CD19CAR-T and BCMACAR-T therapy against B-cell lymphomas, leukemias, and multiple myeloma, a commercial CAR-T product against CD30^+^ hematologic malignancies is still not available. In addition, therapeutic CAR-T cells against CD30^+^ T cell lymphomas present with several challenges, such as the risk of T cell aplasia, fratricide and bidirectional killing, and contamination of CAR-T cell products with malignant T cells [29].

Genome editing approaches are highly innovative and are emerging as a potential replacement for the use of retroviral and lentiviral vectors for T cell engineering [13]. CRISPR/Cas-based gene editing enables precise modification of target genes or insertion of transgenes into cells, and several ongoing preclinical studies promote this technology to develop treatments against cancer, innate genetic defects, and infections. Therefore, we have switched towards the use of non-viral CRISPR/Cas gene editing technology for the production of CAR-T cells [15,16]. Since edited CAR-T cells targeting CD30 may present a novel immunotherapeutic option to control several malignancies, we designed and tested different scFv sequences targeting CD30 and detection domains incorporated into HDRTs for CRISPR-mediated editing. We showed that the improved HDRT incorporating the 5F11 scFv and a tCD34 domain generated high frequencies of CD30CAR^+^ cells with a favorable immunophenotype of naïve and central memory CD30CAR-T cells. The gene-edited cells with physiologically regulated *TRAC*^KI^-CD30CAR expression showed potent cytotoxic properties *in vitro* against several types of CD30^+^ cells (HL, TCL, EBV-LPD). *TRAC*^KI^-CD30CAR-T cells applied five days into NSG-DKO mice after engraftment of L540 cells prevented systemic tumor growth *in vivo* (although some mice showed bioluminescent signals in the anatomical region of the brains). Although significantly higher prolonged survival was observed for CD30CAR-T cell-treated mice, all mice eventually succumbed. At the experimental endpoint, lymphoma cells were detectable at significantly higher frequencies in the bone marrow of control mice compared with CD30CAR-T cell-treated mice, endorsing the therapeutic effect. CD30CAR-T cells were detectable at the endpoint in blood, bone marrow, and spleen, and they were mostly CD4^+^ T cells.

PD-L1/PD-1 blockade antibodies showed impressive capacity to reactivate tumor-infiltrating lymphocytes (TILs) at the tumor microenvironment. Systemic administration of PD-L1/PD-1 blockers promotes major changes in CD8 T cells with clinical antitumor responses. Although some studies supported this hypothesis, it was reported by others that PD-1 knockdown actually reduced the anti-tumor potential of CAR-T cells because it inhibited T cells’ proliferation activity and prevented effector T cells from differentiating into functional memory T cells [30–32]. Thus, in view of further improving the therapeutic effects of CTLs in CD30CAR-T cells, we set up and tested multiplex editing, including *TRAC*^KI^ and PD-1^KO^. We showed that PD-1^KO^ did not negatively affect either the CD8^+^ or the CD4^+^ CD30CAR-T cell expansion and did not compromise CAR expression. However, CD30CAR-PD-1^KO^ T cells did not show superior CTL ability to kill PD-L1^+^ target cells compared to non-PD-1^KO^ controls. Nonetheless, *TRAC*^KI^-CD30CAR-PD1^KO^-T cells showed a trend for higher secretion of cytotoxic proteins (Granzyme A and B) and inflammatory cytokines (IFN-γ and FAS-L) when co-cultured with the lymphoma targets. These results indicated that CD4^+^ Th1 CD30CAR-T cells may have potentially a major effect against the PD-L1^+^ lymphoma targets tested here. Although these results were anticipated, accumulated clinical evidence supports the contribution of CD4 T cells for successful PD-L1/PD-1 blockade immunotherapy [33,34]. In fact, immune responses by circulating CD4^+^ T cells in the peripheral blood of patients treated with PD-1/PDL-1 blockade correlated with clinical responses [35].

As an additional all-in-one strategy for genome editing, we validated a multiplexed gene editing method to knock out PD-1 and/or β2M expression. Compared with *TRAC*^KI^-CD30CAR-T cells, *TRAC*^KI^-CD30CAR-β2M^KO^-T cells showed similar expansion, immunophenotype, and overall cytotoxic activity. T cells lacking expression of HLA-I can greatly facilitate prospective use of third-party or allogeneic CD30CAR-T cells, e.g., to facilitate HLA matching, to avoid CAR-product contamination with malignant T cells, or to enable CAR production when patients with lymphopenia are not eligible due to difficulties in obtaining T cells for CAR-T cell production. Since HLA-II expression was not deleted, *TRAC*^KI^-CD30CAR-β2M^KO^-T cells should be to some extent protected against NK cell cytotoxicity. In comparison to non-HLA-II-expressing target cell lines, it was shown that HLA-DR^+^ target cell lines reduced the cytolytic activity of NK cells, suggesting a “missing-self” hypothesis not only for HLA-I but also HLA-II molecules [36].

A current limitation to further develop combinations of gene knock-ins and knock-outs is the risk of adverse off-target effects, compromising the safety of the edited cells. As described before, simultaneous DSBs can lead to a high rate of genomic rearrangements in edited T cells [37,38]. We observed that CRISPR/Cas9-induced chromosomal translocations were stably maintained over a two-week period in TRAC^KI^-CD30CAR-ß2M^KO^-T cells. These included the expected translocations between the three on-target sites, as well as rearrangements linking these sites to four CAST-Seq–identified off-target loci. At two of these four loci, clear evidence of off-target cleavage was detected, whereas sequence alterations at the remaining two sites did not exceed background levels. At present, it remains unclear whether these latter sites represent false-positive CAST-Seq calls - with its lower limit of detection of approximately 0.01% [20] - or whether true off-target activity occurred but remained below the detection threshold of conventional NGS (∼0.1%). Regardless, the persistence of chromosomal rearrangements raises concerns about potential impacts on genome integrity, therapeutic efficacy, and safety. Careful evaluation and mitigation of these risks will therefore be essential before advancing such CRISPR-edited T cell products into clinical testing.

While several methods and tools are currently available to identify and quantify off-target effects on the genome level, standardized guidelines are still needed to define biological risk predictions [39]. For example, predictive *in vitro* CAR-T cell culture assays and *in vivo* mouse models (such as xenotransplantation into mice reconstituted with the human immune system) could help to evaluate expansion, persistence of immune-functional performance, and risks of genomic rearrangements leading to malignancies. Furthermore, the regulatory bodies may request manufacturing considerations to mitigate all risks of translocations in cell products to be used clinically and traceable data archiving for potential retrospective safety review [39].

One possibility to reduce the risk of DSBs during multiplex editing is to exploit CRISPR base editors, enabling targeted mutagenesis with limited translocations [40].For all-in-one TRAC-KI and additional knock-outs, orthogonal CRISPR enzymes for nuclease-driven knock-in and base editing mediated knockouts can be used to prevent translocations. For instance, Glaser et al 2023 combined CRISPR-Cas12 Ultra nuclease-assisted knock-in and Cas9-derived base editing technology for knock-outs within a single intervention [41]. They showed efficient TRAC^KI^-CD19CAR, along with two knockouts that silenced the expression of the major histocompatibility complexes (MHC) class I and II. Recently, Glaser et al and Skeate et al. further simplified multiplex knockout and simultaneous knock-in by repurposing Cas9-derived base editors for knock-in using just a single enzyme [42,43]. Several groups demonstrated that both strategies (orthogonal Cas enzymes or base editors only) reduced balanced translocations between the on-target sites to <0.05% of edited CAR-T cells [42–45].

To effectively translate this strategy into clinical application, an orthogonal approach should be adopted - incorporating advanced in silico prediction models and complementary in vitro methods to identify editors that combine high activity with high specificity, followed by CAST-Seq analyses in edited T cells to identify off-target sites and evaluate genome stability in the clinically relevant cell type. This integrated framework would not only enhance the detection and quantification of off-target effects but also improve the reliability and safety assessment of gene editing outcomes prior to clinical implementation.

## Conclusion

In conclusion, this work advanced towards new generations of potent gene-edited *TRAC*^KI^-CD30CAR-T cells that no longer rely on viral vectors for manufacturing. CD30CAR-T cells generated with lentiviral vectors and incorporating the 5F11 scFV have been tested clinically and showed transient responses and some signs of toxicity, and therefore the *TRAC*^KI^-CD30CAR-T cells can potentially have superior persistency and safety advantages due to the putative physiological control of the CAR expression by the *TRAC* promoter [46]. We were able to demonstrate feasible generation of all-in-one *TRAC*^KI^-CD30CAR-PD1^KO^-T cells, which demonstrated the relevance of the CD4^+^ subpopulation to secrete cytokines in response to tumor exposure. We also validated *TRAC*^KI^-CD30CAR-β2M^KO^-T cells in terms of feasibility and maintained potency *in vitro*. The generation of the triple KO combination *TRAC*^KI^-CD30CAR-PD1^KO^-β2M^KO^-T cells was in principle feasible; however, their expansion was affected, and several translocations were observed. These new gene editing methods will now require upscaling, GMP adaptations, and critical analyses of genomic alterations for translation of these “next generation” cell therapies into treatment of patients with advanced CD30^+^ pathologies.

## List of Abbreviations

β2M: Beta 2 microglobulin
CAR: Chimeric antigen receptor
CRISPR/Cas: Clustered regularly interspaced short palindromic repeats/Cas
EBV: Epstein-Barr virus
EBV-LPD: Epstein–Barr virus–associated lymphoproliferative diseases
FACS: Fluorescence-activated cell sorting
HIV: Human Immunodeficiency virus
HLA: Human leukocyte antigen
GMP: Good Manufacturing Practice
KI: Knock-in
KO: Knock-out
LCL: Lymphoblastoid cell line
PCR: Polymerase chain reaction
PD-1: Programmed cell death protein 1
PD-L1: Programmed cell death 1 ligand 1
TCL: T-cell lymphoma
TCR: T-cell receptor
TRAC: T-cell receptor alpha chain
WT: Wild-type

## Declarations

## Regulatory approvals

Leukapheresis units were obtained by the Institute of Transfusion Medicine of the Hannover Medical School in accordance with ethics study protocols approved by the Ethics Review Board (approval Nr. 4837) and the University Hospital Cologne (approval Nr. 22-1423_3). Peripheral blood samples (“buffy coats”) were obtained by the Institute of Transfusion Medicine under written informed consent of the donors in accordance with study protocols approved by University Hospital Cologne Ethics Review Board (approval Nr 23-1309). All animal experiments have been approved by LAVE (Landesamt für Verbraucherschutz und Ernährung) approval number Az 81-02.04.2023.A155.

## Consent for publication

Not applicable.

## Availability of data and material

All data relevant to the study are included in the article and in the uploaded supplement information. Additional data is available upon request to the communicating author.

## Author’s Email contacts and contributions

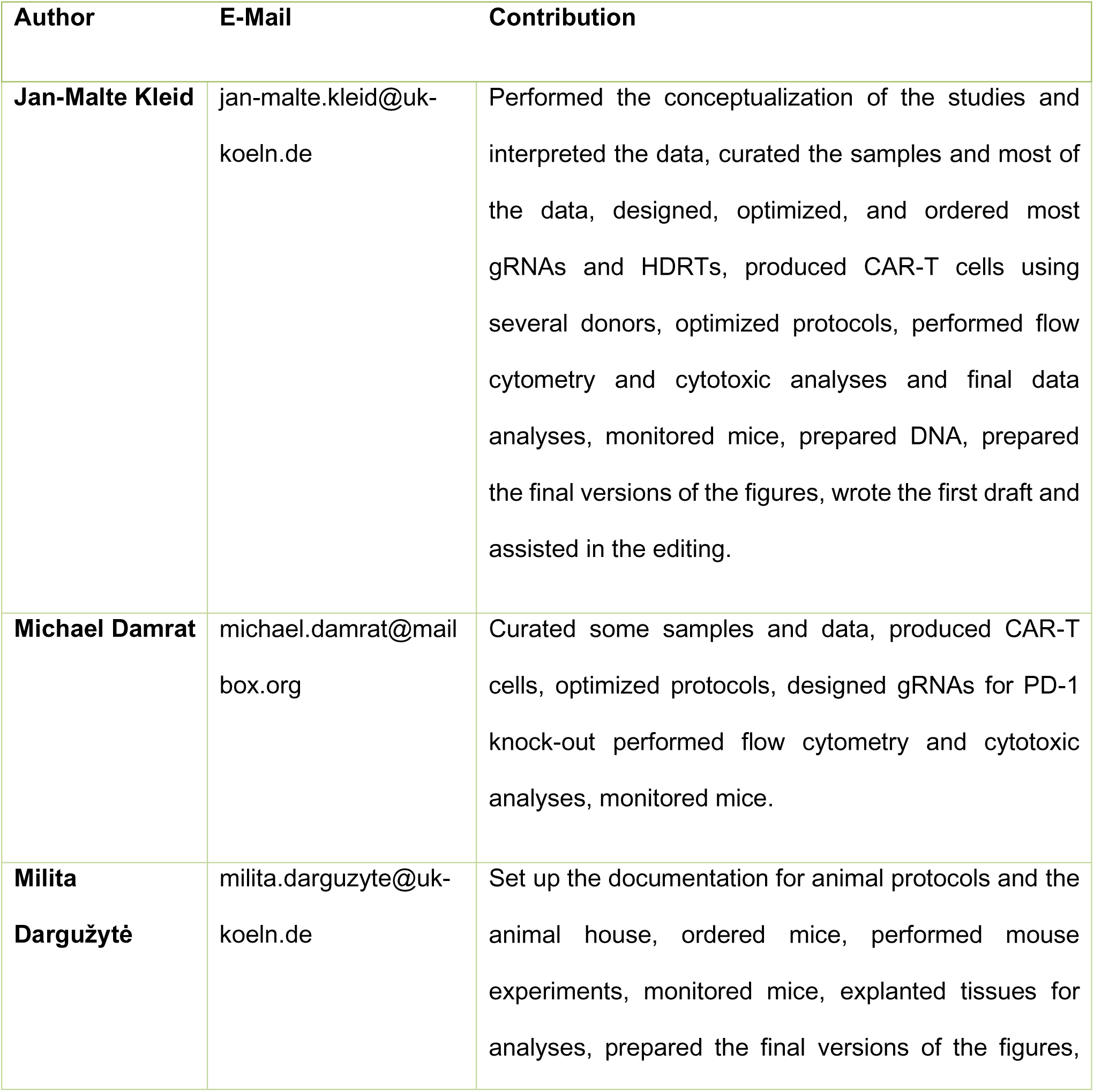
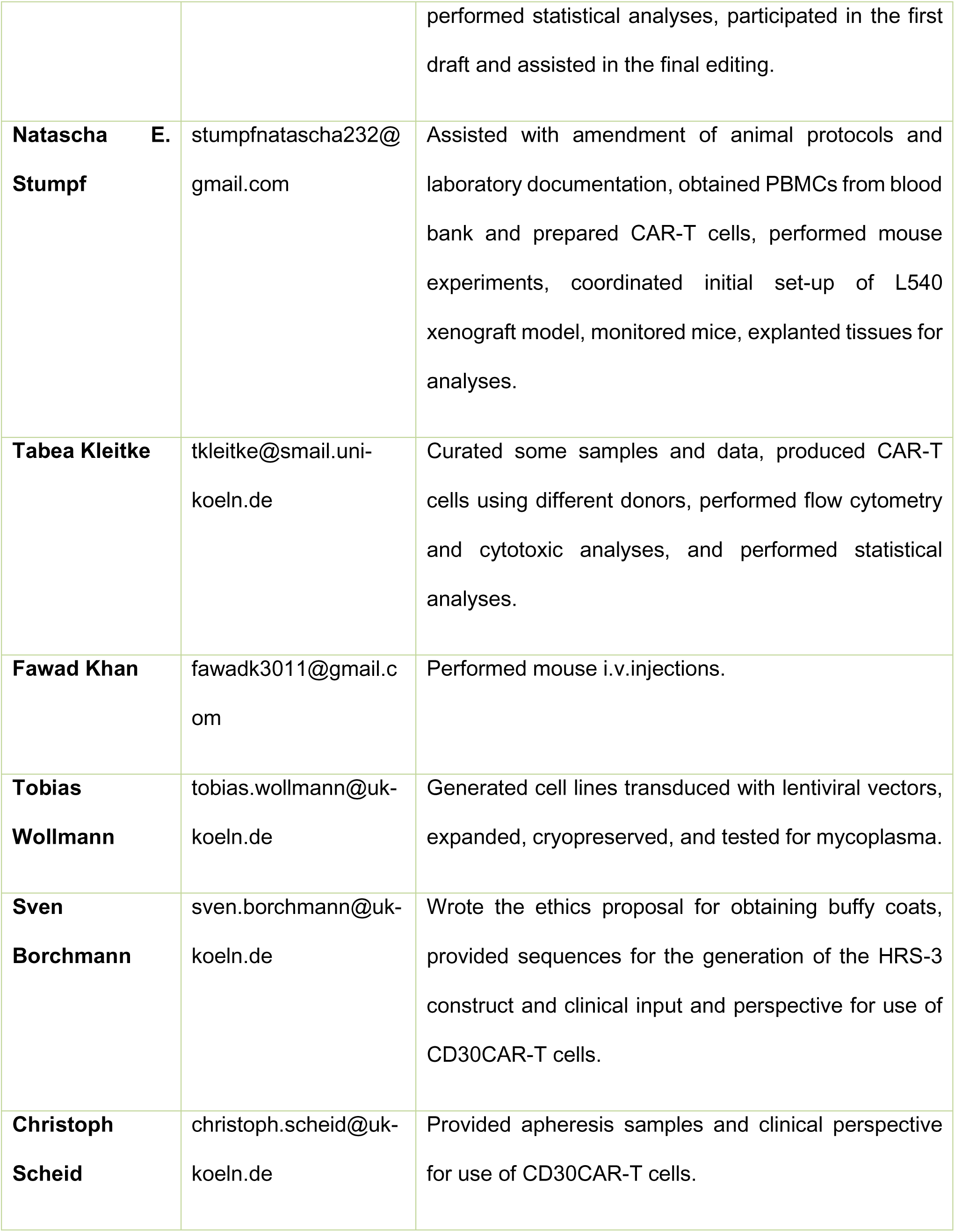
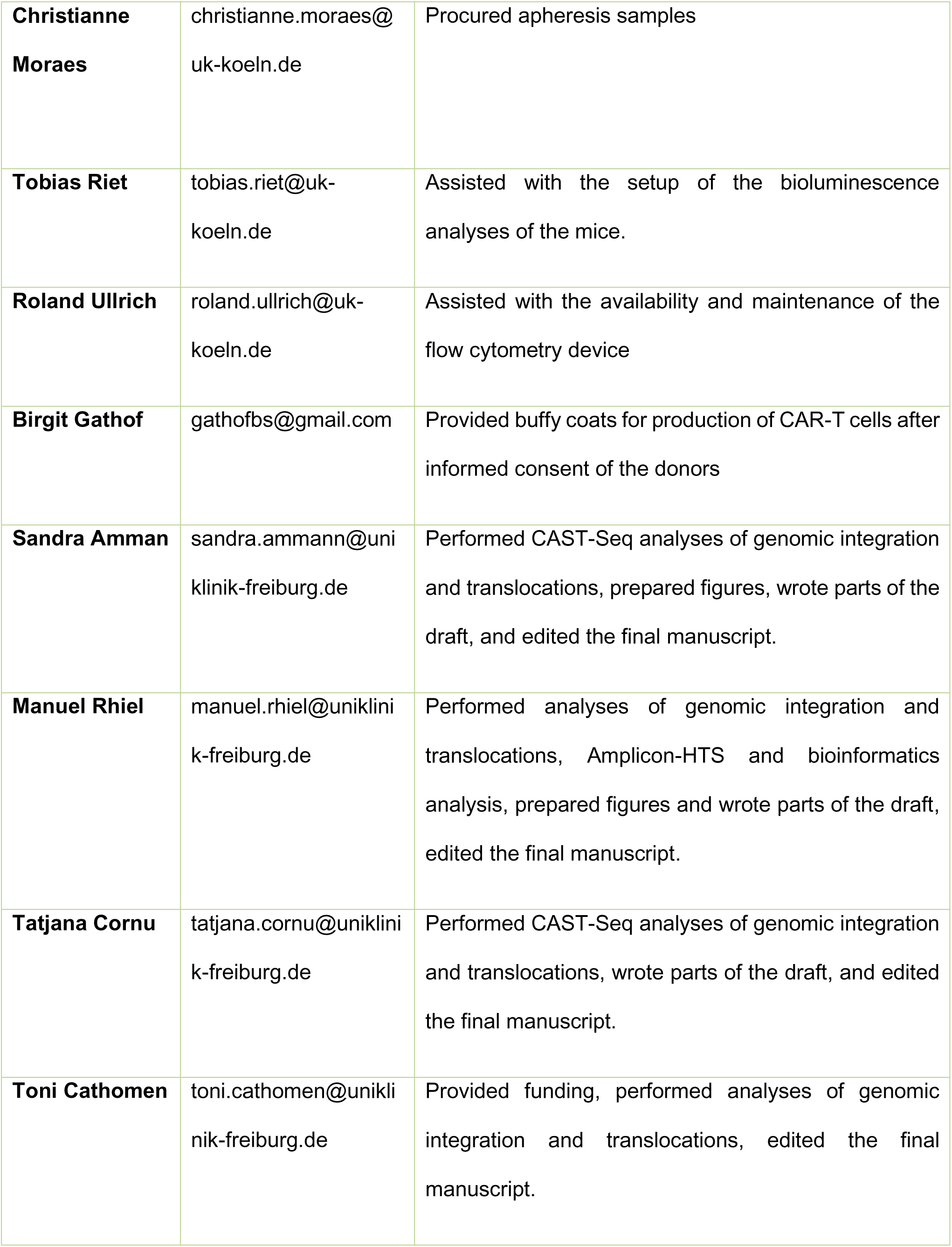
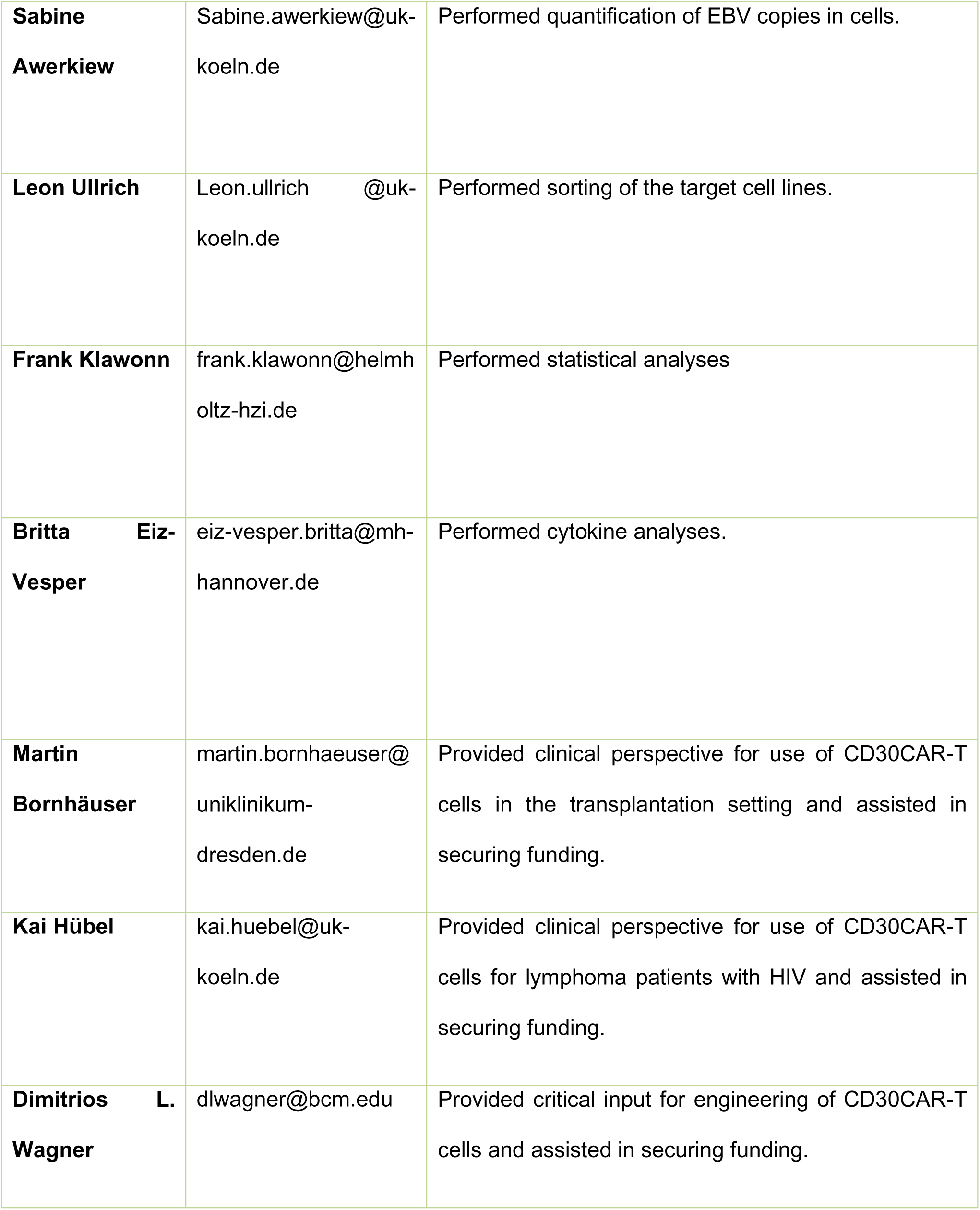
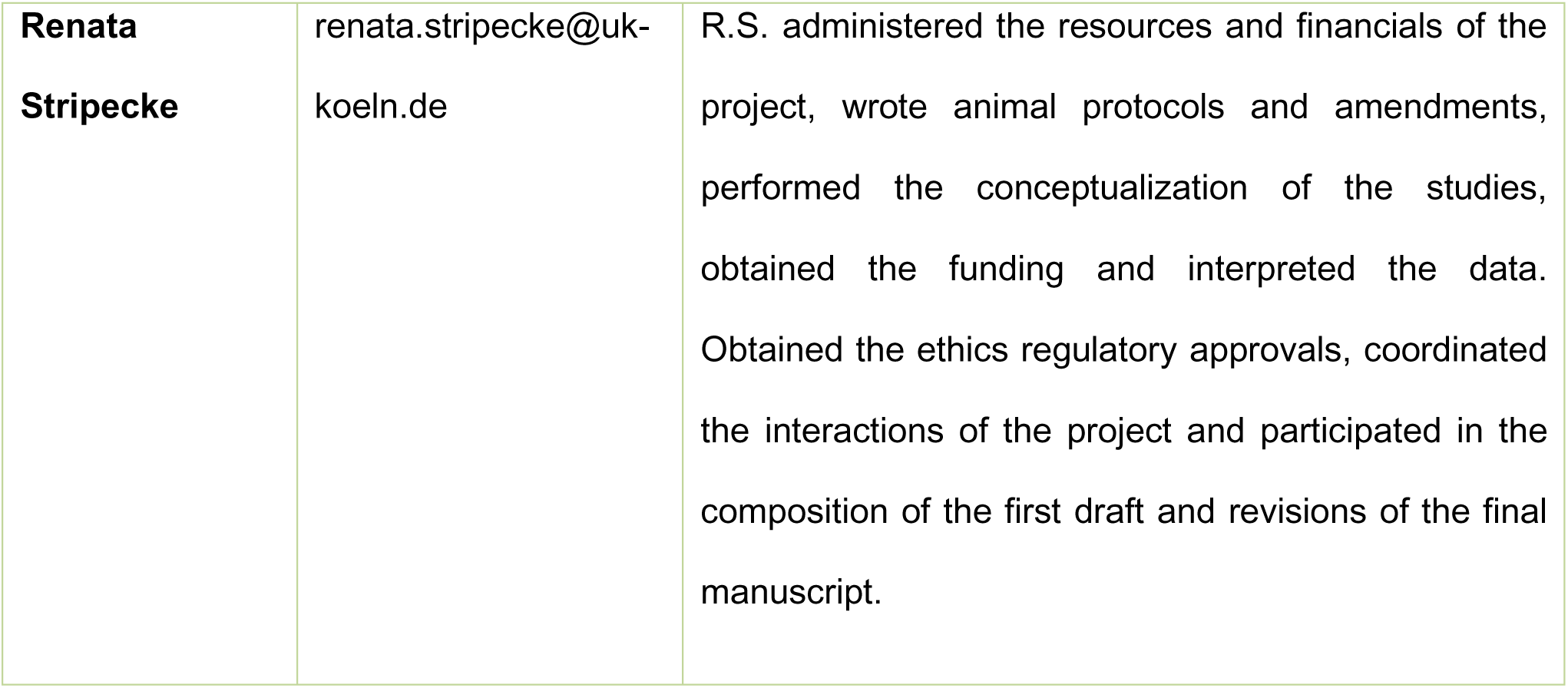

**All authors have read and agreed to the published version of the manuscript.**

## Acknowledgements

The authors thank colleagues of the Institute of Translational Immuno-Oncology and Clinical Department of Internal Medicine-I and all the staff of the Institute of Transfusion Medicine of the University Hospital Cologne who supported our work. We thank all the staff of the animal house of the University Hospital Cologne for the important assistance with preparation of animal protocols and keeping of the mice. We thank Prof. Axel Schambach for the lentiviral construct to mark the lymphoma lines. We thank Geoffroy Andrieux for the bioinformatic analysis of CAST-Seq samples.

## Additional files

| File name | File format | Title of data | Description |
| --- | --- | --- | --- |
| **Supplement Table 1** | Microsoft Word (docx) | Antibodies | List of antibodies used for flow cytometry |
| **Supplement Table 2** | Microsoft Excel spreadsheet (xlsx) | Primers | List of primers used for CAST-seq |
| **Supplement Table 3** | Microsoft Excel spreadsheet (xlsx) | Translocations | Quantification of translocations detected by CAST-seq |

