## Supplementary figures and images for "Next-generation all-in-one CRISPR/Cas9 multiply-edited CD30CAR-T cells: Potency despite risk of translocations"

### Supplement Figure S1

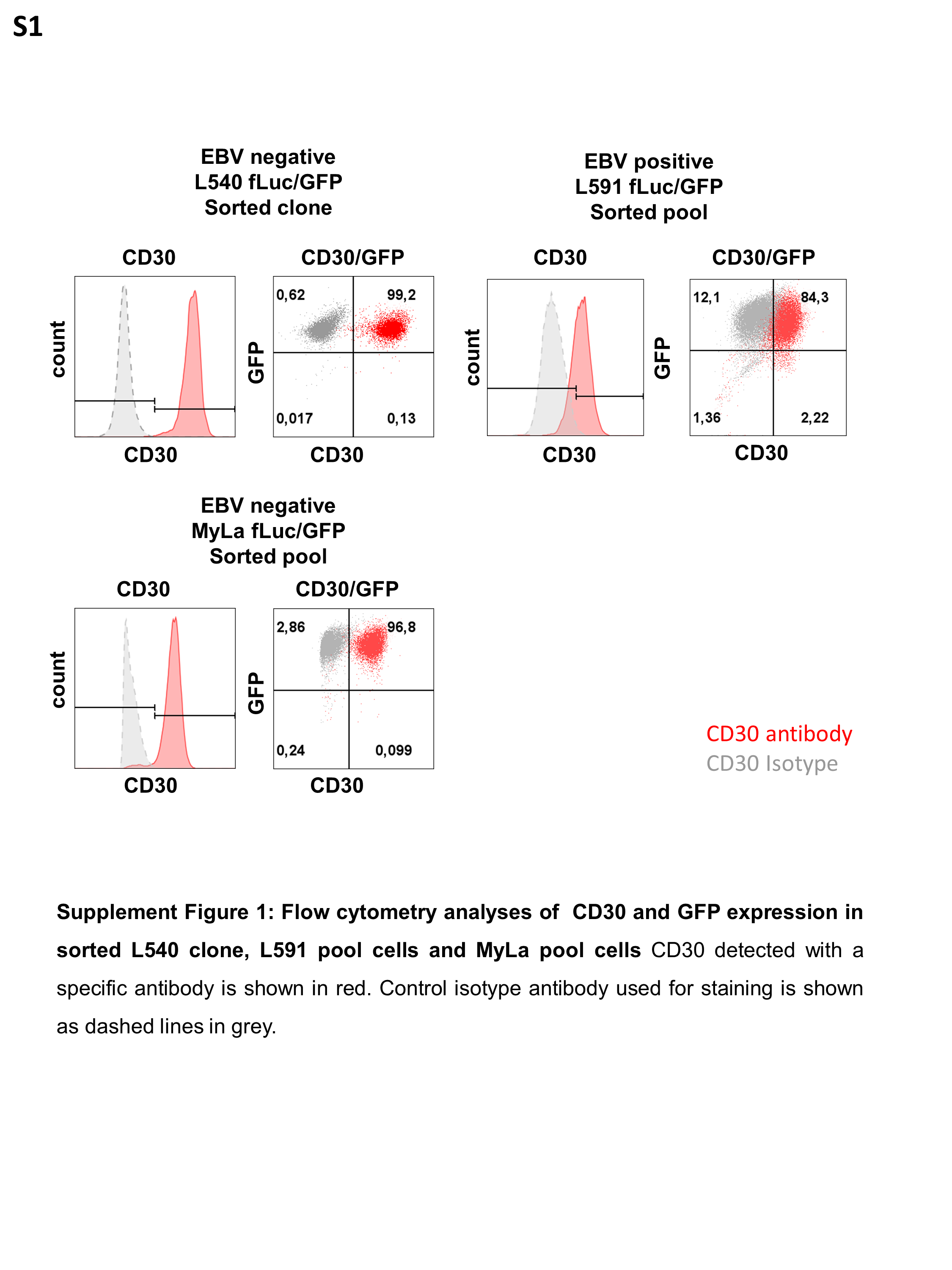

### Supplement Figure S2

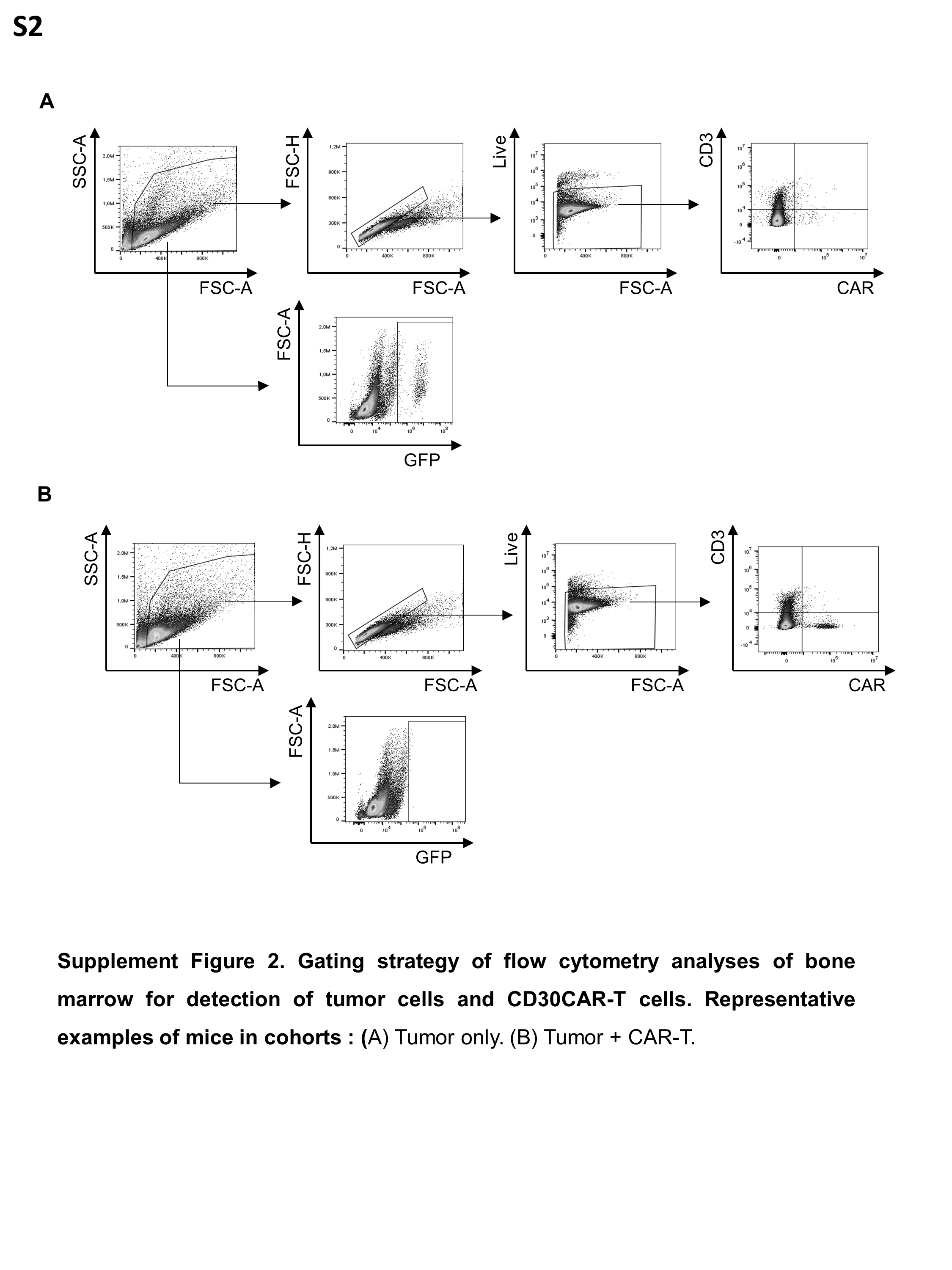

### Supplement Figure S3

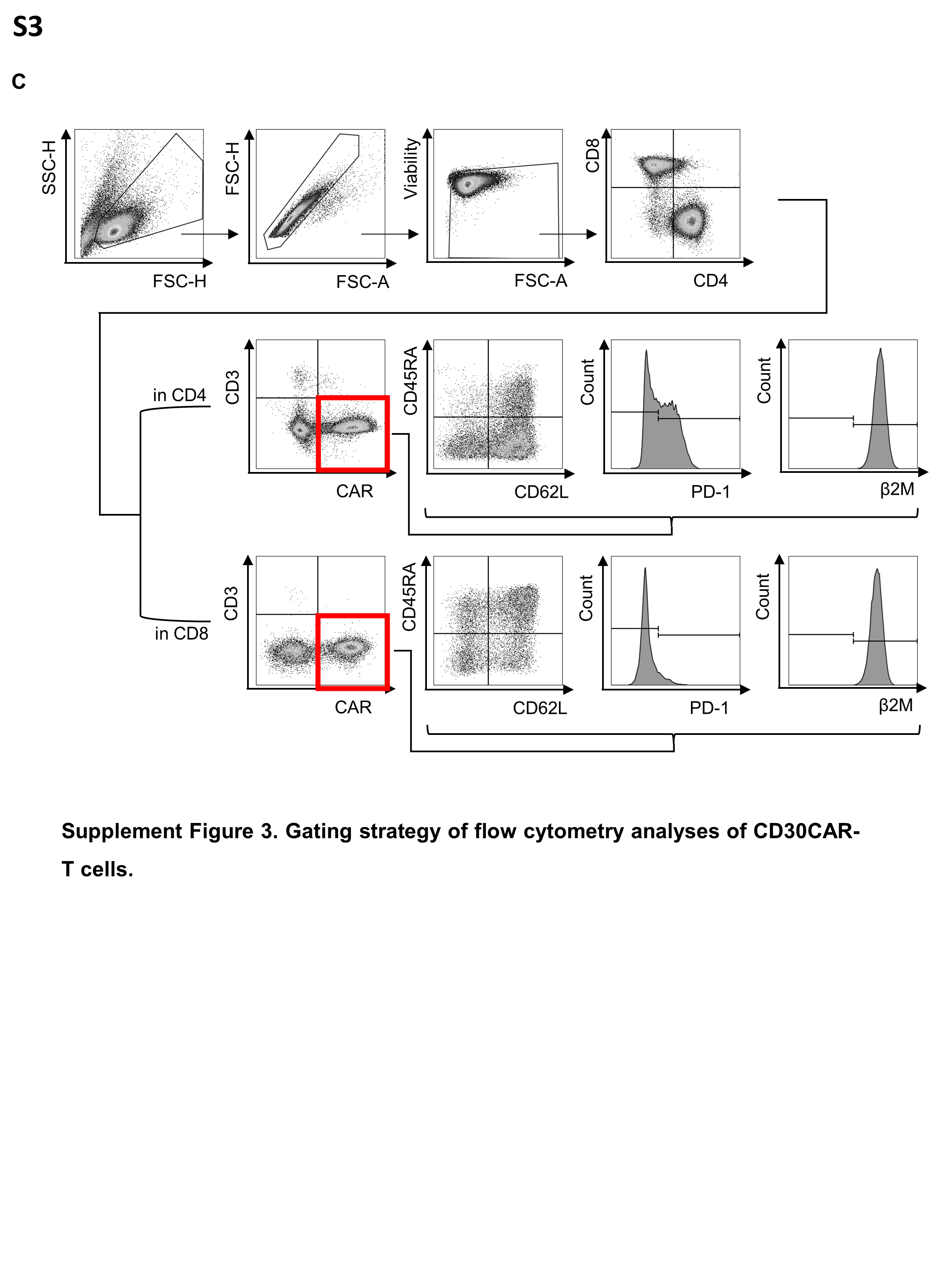

### Supplement Figure S4

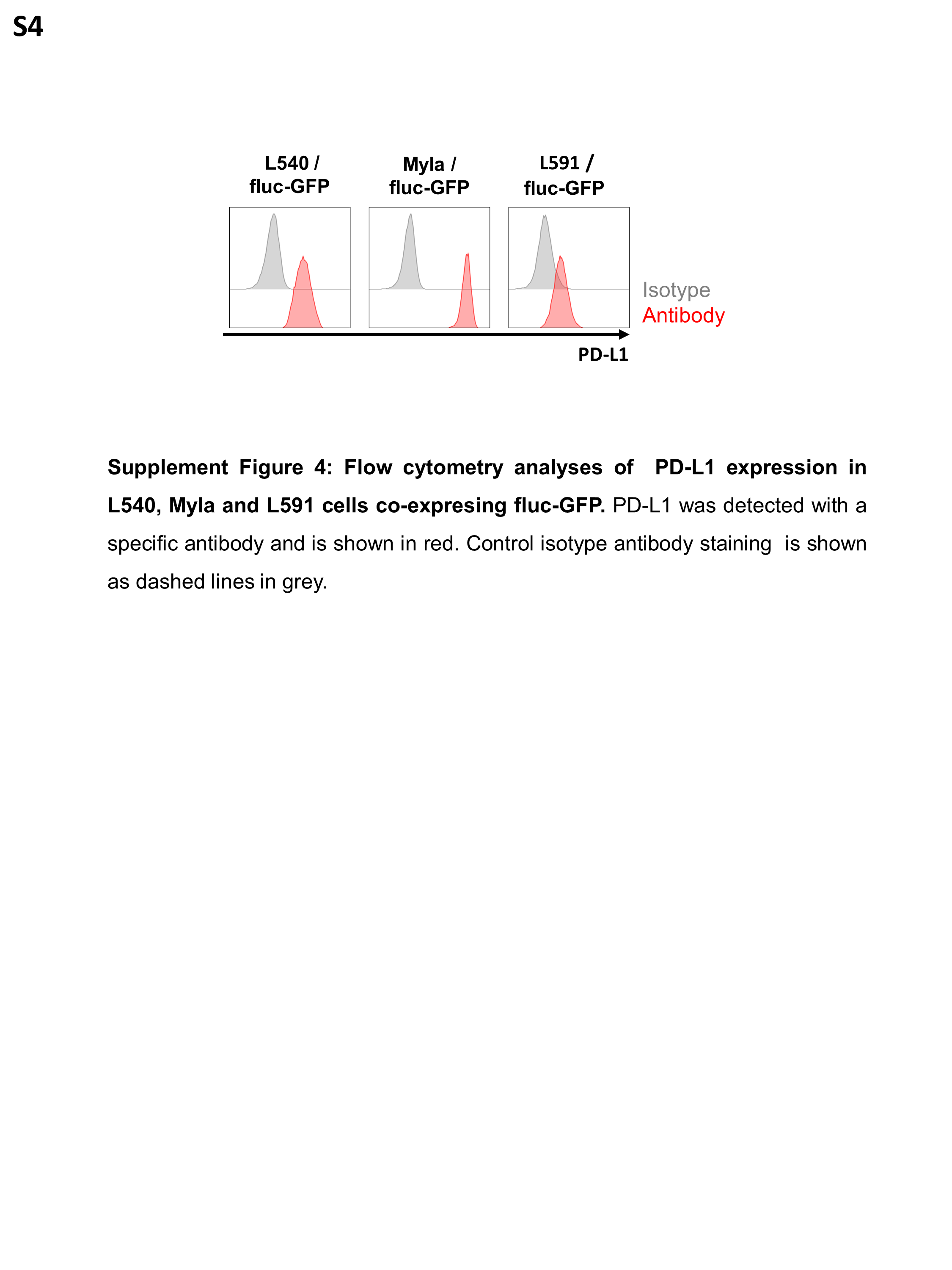

### Supplement Figure S5

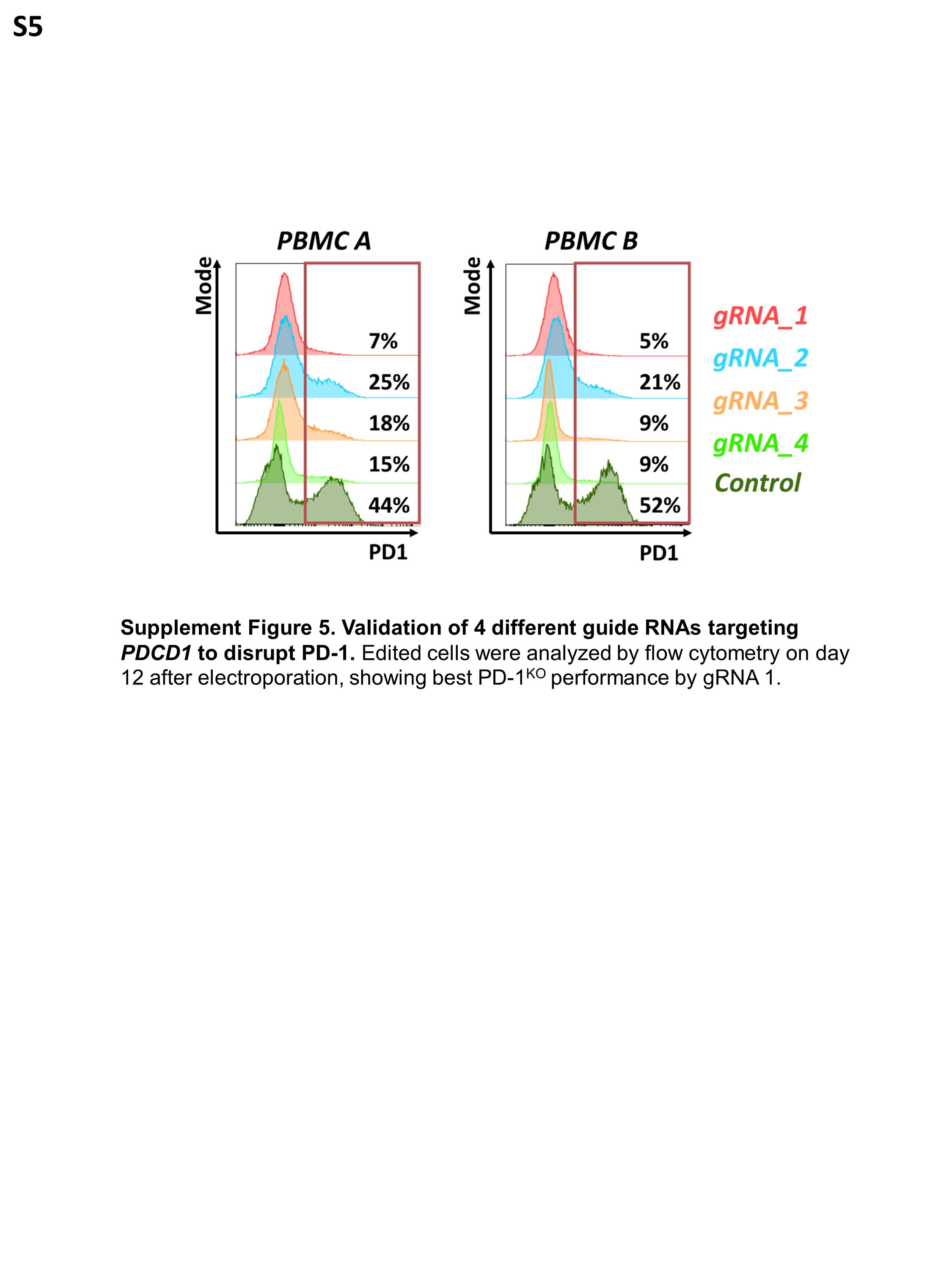

### Supplement Figure S6

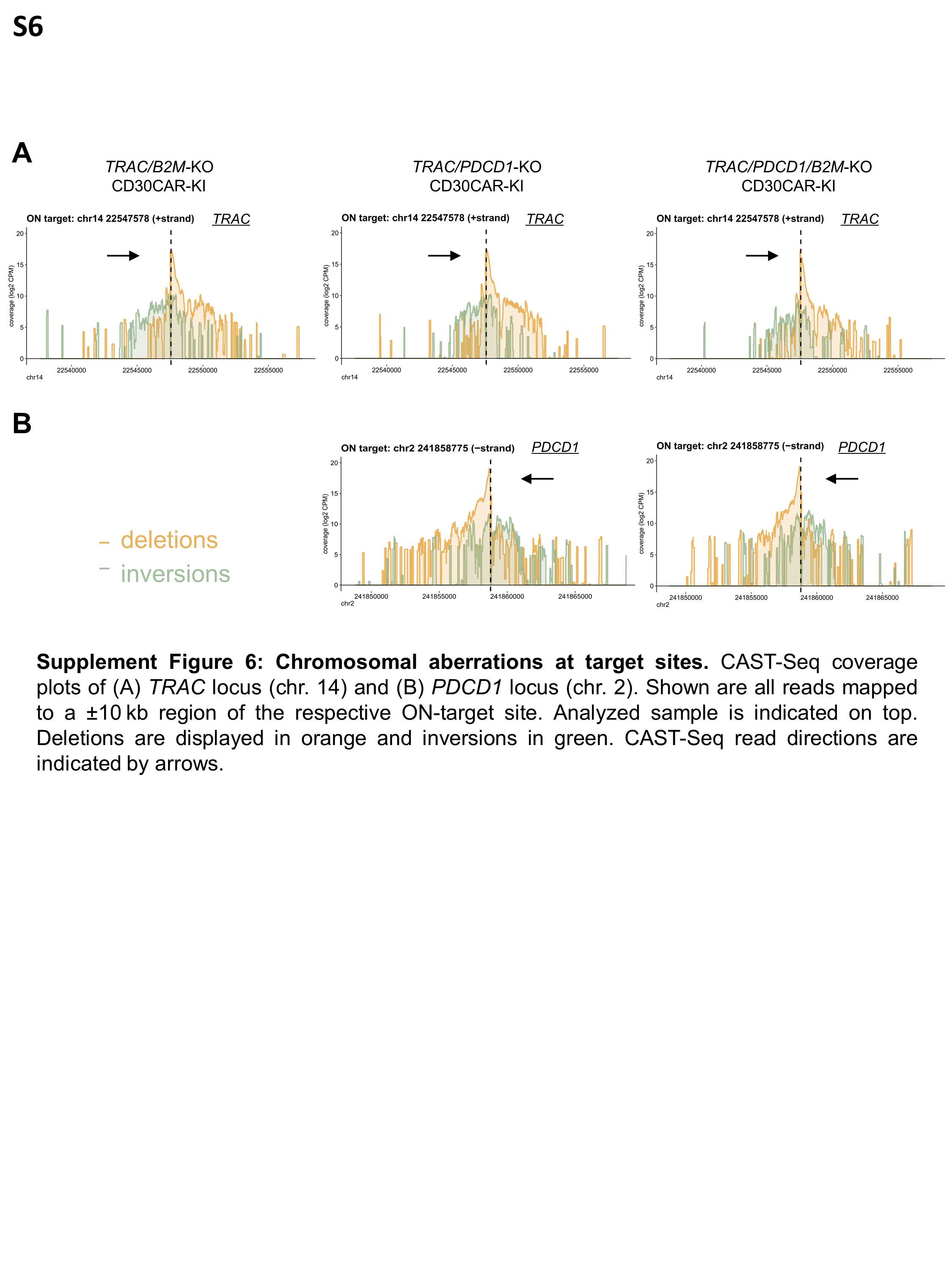
