## Supplement Table 1 for "Next-generation all-in-one CRISPR/Cas9 multiply-edited CD30CAR-T cells: Potency despite risk of translocations"

| Target | Fluorophore | Clone | Vendor | Cat Nr. | Dilution |
| --- | --- | --- | --- | --- | --- |
| Hu IgG1 for CAR detection | Alexa Fluor 647 | Polyclonal | Jackson Immuno Research Laboratories | 109-606-170 | 1:100 |
| CD34 | Alexa Fluor 647 | QBEND10 | BD | 568772 | 1:100 |
| Viability | eFluor450 (BV421) |  | invitrogen | 65-0863-14 | 1:1000 |
| CD3 | Pe-Dazzle594 | HIT3a | Biolegend | 300335 | 1:100 |
| CD4 | PerCP | OKT4 | Biolegend | 317432 | 1:400 |
| CD8 | BV650 | RPA-T8 | Biolegend | 301042 | 1:200 |
| PD1 | PE | EH12.2H7 | Biolegend | 329906 | 1:100 |
| HLA A, B, C | APC-Cy7 | W6/32 | Biolegend | 311426 | 1:100 |
| CD45RA | BV 605 | HI100 | Biolegend | 304134 | 1:75 |
| CD62L | Pe-Cy7 | DREG-56 | Biolegend | 304821 | 1:50 |
| CD30 | BV510 | BerH8 | BD | 744407 | 1:100 |
| PD-L1 | APC | 29E2A2 | Biolegend | 329708 | 1:100 |

Supplement Table 1: List of antibodies used for flow cytometry analyses.
